# Machine-learning-based predictions of caloric restriction associations across ageing-related genes

**DOI:** 10.1101/2021.07.17.452785

**Authors:** Gustavo Daniel Vega-Magdaleno, Vladislav Bespalov, Yalin Zheng, Alex A. Freitas, Joao Pedro de Magalhaes

## Abstract

Caloric restriction (CR) is the most studied pro-longevity intervention; however, a complete understanding of its underlying mechanisms remains elusive, and new research directions may emerge from the identification of novel CR-related genes and CR-related genetic features. This work used a Machine Learning (ML) approach to classify ageing-related genes as CR-related or NotCR-related using 9 different types of predictive features: PathDIP pathways, two types of features based on KEGG pathways, two types of Protein-Protein Interactions (PPI) features, Gene Ontology (GO) terms, Genotype-Tissue Expression (GTEx) expression features, GeneFriends co-expression features and protein sequence descriptors. Our findings suggested that features biased towards curated knowledge (*i.e*. GO terms and biological pathways), had the greatest predictive power, while unbiased features (mainly gene expression and co-expression data) have the least predictive power. Moreover, a combination of all the feature types diminished the predictive power compared to predictions based on curated knowledge. Feature importance analysis on the two most predictive classifiers mostly corroborated existing knowledge and supported recent findings linking CR to the Nuclear Factor Erythroid 2-Related Factor 2 (NRF2) signalling pathway and G protein-coupled receptors (GPCR). We then used the two strongest combinations of feature type and ML algorithm to predict CR-relatedness among ageing-related genes currently lacking CR-related annotations in the data, resulting in a set of promising candidate CR-related genes (*GOT2, GOT1, TSC1, CTH, GCLM, IRS2* and *SESN2*) whose predicted CR-relatedness remain to be validated in future wet-lab experiments.

## 1 Introduction

Ageing increases the risk of disease and death as it declines homeostasis and decreases the capacity to respond to environmental stimuli (MacNee, 2016). Given the widespread interest in reversing and ultimately preventing the detrimental effects of ageing, considerable effort has been devoted to understanding its underlying biochemical mechanisms (de Magalhães et al., 2012). It is known that ageing-related changes are multifactorial and involve a variety of processes, including genomic instability, telomere attrition, epigenetic alterations, loss of proteostasis, impaired nutrient sensing, mitochondrial dysfunction, cellular senescence, stem cell exhaustion, and altered intracellular communication (López-Otín et al., 2013).

Caloric Restriction (CR), which involves reducing total dietary energy intake while maintaining adequate vitamin and mineral levels, is currently the most promising intervention for increasing both lifespan and healthspan, as experiments with a variety of species have shown that CR not only induces longevity but also retards the ageing process (de Magalhães et al., 2012). Although the mechanism underlying these pro-longevity effects is unknown, evidence suggests that CR: 1) reduces oxidative damage by reducing the production of Reactive Oxygen Species (ROS); 2) decreases circulating insulin and glucose levels, resulting in decreased cell growth and division and a shift toward maintenance and repair; and 3) decreases growth hormone and insulin-like growth factor levels (Gillespie et al., 2016).

Additionally, a wealth of publicly available omics data on ageing has emerged thanks to new high-throughput sequencing technologies (Wieser et al., 2011). Hence, a relatively recent approach for studying the ageing process is based on Machine Learning (ML) techniques that learn patterns about gene or protein functions by analyzing gene or protein features such as Gene Ontology (GO) terms, metabolic pathways, and protein-protein interactions, to name a few (Fabris et al., 2017). Examples include the association of human genes with ageing-related diseases (Fabris et al., 2020); prediction of gene deletion effects on yeast longevity (Huang et al., 2012); and the determination of blood age (Weidner et al., 2014); among others (Zhavoronkov et al., 2019).

This work aims to identify novel CR-related candidate genes from ageing-related genes while also identifying genetic features which increase the likelihood that certain ageing-related genes become CR-related. To accomplish this, we created 11 datasets based on 9 different types of predictive features and two approaches to combine all those features into an integrated dataset. The 9 used feature types were: PathDIP pathways, two types of features based on KEGG pathways, two types of Protein-Protein Interactions (PPI) features, Gene Ontology (GO) terms, Genotype-Tissue Expression (GTEx) expression features, GeneFriends co-expression features and protein sequence descriptors. These datasets provide a wealth of information for representing each of the ageing-related genes under study (*i.e*., genes retrieved from the GenAge database, as explained in Section 2.1.1). Then, a ML approach was used to predict whether or not an ageing-related gene is CR-related (*i.e*., using information from the GenDR database of CR-related genes, as explained in Section 2.1.1). The approach involved comparing the predictive accuracy of four different tree-based ensemble ML algorithms across all 11 datasets, with the most predictive ML algorithm, Dataset combinations being then used for feature importance analysis, which led to the identification of key CR-related genetic attributes. Finally, we used the two best performing classifiers to infer potential under-explored CR-relatedness from ageing-related genes lacking CR-related annotations in the dataset.

Our findings indicate that the most predictive features are based on curated knowledge, such as GO terms and biological pathways. The least predictive features were gene expression and co-expression features. Apart from the well-established CR-related autophagy and longevity regulating pathways, our findings indicate that the G Protein-Coupled Receptor (GPCR) signalling, cellular responses to external stimuli, and Nuclear Factor Erythroid 2-Related Factor 2 (NRF2) pathways were among the most significant features for inferring CR-relatedness. This is consistent with recent evidence indicating that these pathways act as mediators of CR effects. Additionally, the oxidation-reduction reaction was the most predictive feature across all GO terms. Finally, predictions from the strongest classifiers indicate that the ageing-related genes *GOT2, GOT1, TSC1, CTH, GCLM, IRS2*, and *SENS2* may share an under-explored association with CR.

## 2 Methods

### 2.1 Datasets construction

This and the following sections use the following ML terminology: an instance refers to any ageing-related gene/protein included in the datasets; whereas a feature is any observable property or attribute of any instance (*e.g*., GO term annotations, association with biological pathways, protein sequence descriptors, etc).

#### 2.1.1 Ageing genes and CR labels retrieval

GenAge (Tacutu et al., 2018) is a benchmark database of ageing-related genes. GenAge “model organisms” is a subsection of GenAge consisting of genes in model organisms that, if genetically modulated, result in significant changes in the ageing phenotype and/or longevity (de Magalhaes et al., 2009). The majority of observations have been made on mice, nematodes, fruit flies and budding yeast. In this work, ageing-related genes from those four organisms were downloaded from GenAge Build 20 (09/02/2020). Then, human orthologs of these genes were retrieved from the OMA Orthology database (Train et al., 2017) (2020 Release) using the OMA browser’s Genome Pair View, which allows for the download of orthologs between two species (https://omabrowser.org/oma/genomePW/). Since some genes in different organisms are mapped to overlapping human or-thologs, only one gene from each set of repeated genes was retained, resulting in 1137 human ageing-related genes.

GenDR (Wuttke et al., 2012) is a database of Dietary Restriction (DR)-related genes. DR-essential genes are defined in GenDR as those which, if genetically modified, impair DR-mediated lifespan extension. In this work, 215 CR-associated genes from the aforementioned four model organisms were retrieved from the “Gene Manipulation” section of the GenDR database, build 4 (24/06/2017), and used as input for another OMA human orthologs query, which led to 152 human CR genes after keeping only one of each repeated ortholog gene coming from different organisms. The ageing- and CR-related human genes retrieval processes are summarised in Figure 1.A.

**Figure 1:**
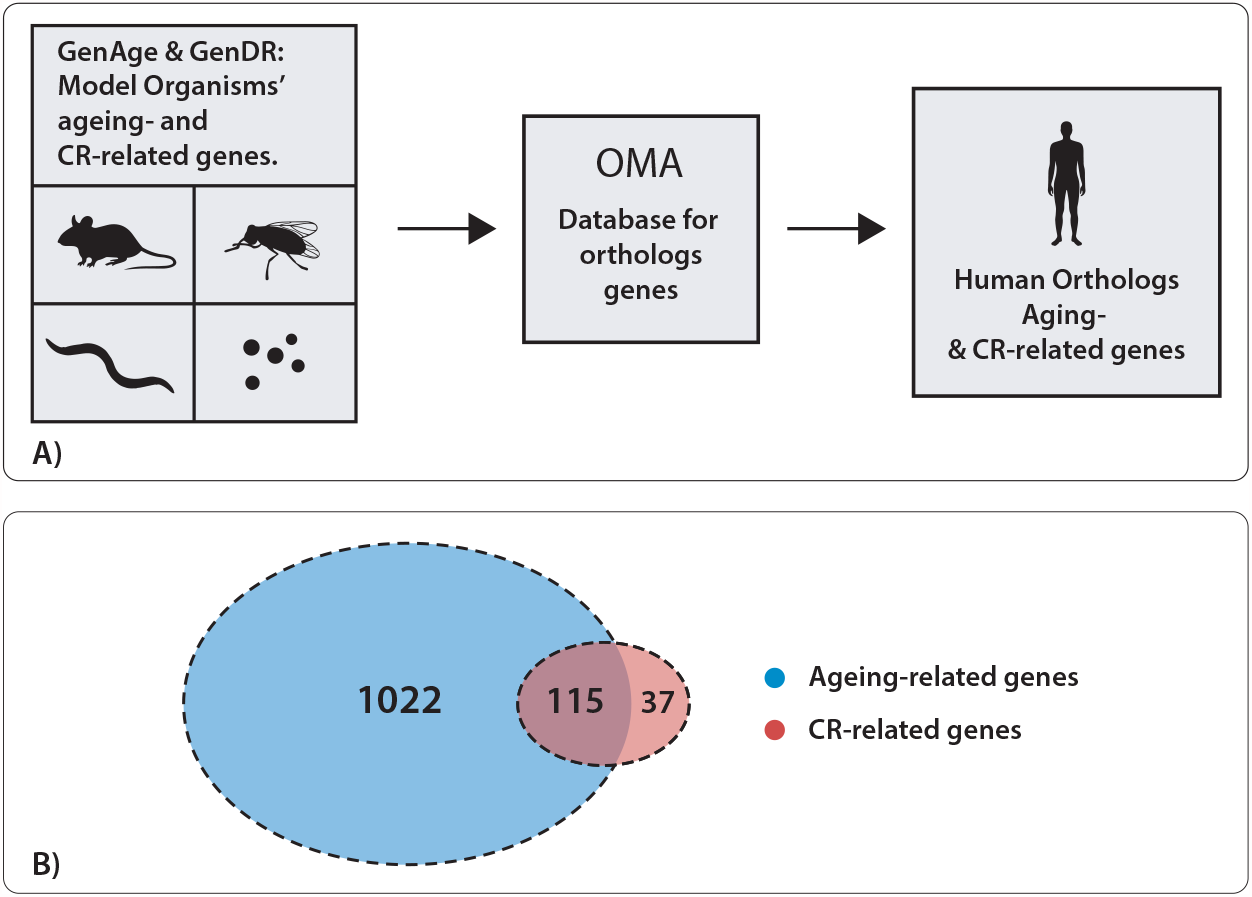
**A**. Ageing- and CR-related genes from mice, fruit fles, nematodes, and budding yeast were retrieved from GenAge and GenDR. Then, an OMA database query was performed to retrieve their corresponding human orthologs, which are the ageing-related and CR-related genes used in this work. **B**. Human ageing- and CR-related genes’ overlap.

Finally, the overlap of the retrieved ageing- and CR-related human orthologs resulted in 115 genes (Figure 1.B) which were labelled as *Ageing*_*CR*_-related, while the remaining 1022 ageing-related genes that didn’t overlap with CR-related orthologs were labelled as *Ageing*_*NotCR*_-related.

#### 2.1.2 PathDIP Dataset

This dataset consists of binary features, *i.e*., each feature value indicates whether or not an ageing-related gene belongs to corresponding specific PathDIP pathway. To accomplish this, we queried the PathDIP (Rahmati et al., 2017) database (version 4.0.7.0) to download a dataset in which instances were ageing-related genes, and features were biological pathways coming from a variety of database sources, including ACSN2, BioCarta, EHMN, HumanCyc, INOH, IPAVS KEGG, NetPath, OntoCancro Panther Pathway, PharmGKB, PID, RB-Pathways, REACTOME, stke systems-biology.org, SignaLink2.0, SIGNOR2.0, SMPDB, Spike, UniProt Pathways, and WikiPathway. The resulting dataset contained 1640 pathways and 986 ageing-related genes: 110 labelled as *Ageing*_*CR*_-related and 876 labelled as *Ageing*_*NotCR*_-related.

#### 2.1.3 KEGG-Pertinence Dataset

This dataset consists of binary features, where each feature indicates whether or not an ageing-related gene (instance) belongs to a specific KEGG pathway. To construct this dataset, a data frame relating KEGG pathways with the genes they contain was retrieved by using the *getGeneKEGGLinks* command of the *R*’s Lima package (Ritchie et al., 2015). Then, only the KEGG pathways that were associated with at least one of the ageing-related genes were retained, yielding 312 KEGG pathways. The resulting dataset contained 799 ageing-related genes: 94 labelled as *Ageing*_*CR*_-related and 705 labelled as *Ageing*_*NotCR*_-related.

#### 2.1.4 KEGG-Influence Dataset

This approach was inspired by the feature-creation method proposed in (Fabris and Freitas, 2016). Instead of producing binary features like KEGG pertinence, that method examines the internal contents of each KEGG pathway to produce numerical features, where each feature value measures the extent to which each protein influenced all the other proteins in the pathway. Figure 2.A illustrates the influence that a reference protein (red node in the graph) exerts on other proteins (the remaining graph’s nodes) within a given pathway. In essence, proteins coloured in dark blue can be “reached” only via upstream paths passing through the reference protein and thus are highly influenced by it. As the proteins in the pathway can be reached via more upstream paths not involving the reference protein, they become less influenced by it, and are represented by lighter blue colours. Proteins that receive no influence are coloured in white. Supplementary materials (S.1.1) contain a complete description of this dataset which contains 1770 features and 799 ageing-related genes, 94 of which are labelled as *Ageing*_*CR*_-related and 705 labelled as *Ageing*_*NotCR*_-related.

**Figure 2:**
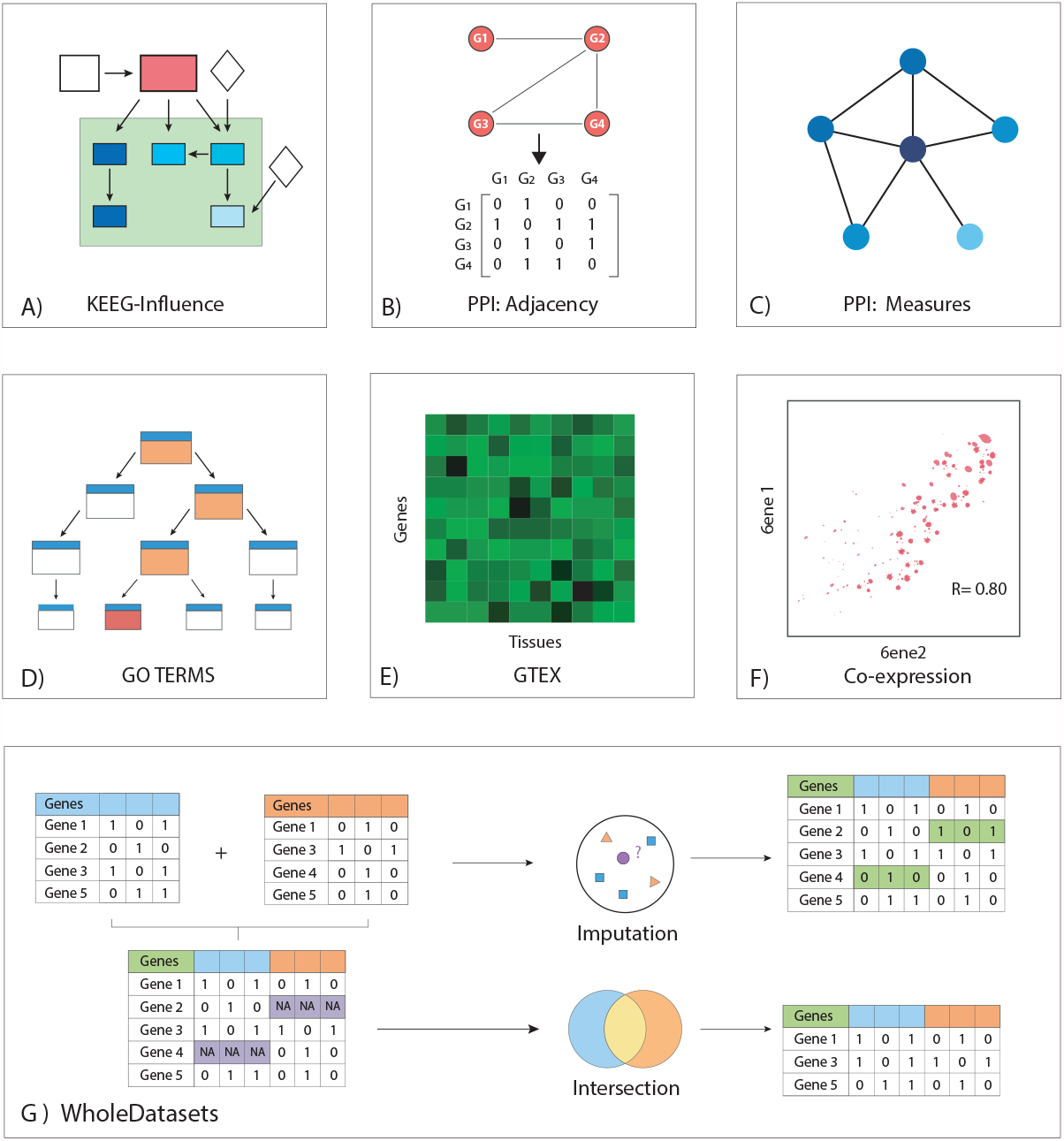
Graphical overview of the datasets used in this work. **A**. KEGG-Influence. **B**. PPI Adjacency. **C**. PPI measures. **D**. GO terms. **E**. GTEx. **F**. Co-expression **G**. WholeDatasets.

#### 2.1.5 Protein-Protein Interaction (PPI) Adjacency Dataset

The human physical PPI network was downloaded from *BioGrid* (Release 3.5.185): *BIOGRID-MV-Physical-3.5.181.tab2.zip* (Stark et al., 2006). *Interactions were then processed using the R*’s igraph library to create a graph object with a total of 25,292 gene products and 324,892 interactions. After removing loops and repeated edges, the graph consisted of 25,292 nodes and 92,237 edges. This graph contained 850 ageing-related genes, 86 of which were labelled as *Ageing*_*CR*_-related while the remaining 764 as *Ageing*_*NotCR*_-related. Then, a dataset with binary features was extracted from the PPI-graph’s adjacency matrix. The *ij*-*th* element of this matrix takes the value 1 if the gene-products *G*_*i*_ and *G*_*j*_ are adjacent in the PPI-graph (*i.e*., if there is an edge connecting them), or 0 otherwise. We retained only ageing-related genes of the adjacency matrix as instances, whereas for features, we kept not only ageing-related genes but also genes not associated with ageing that directly interact with at least one of the ageing-related genes, yielding 5718 features. Figure 2.B depicts a representation of this dataset where an adjacency matrix is constructed from a graph.

#### 2.1.6 PPI Graph Measures Dataset

This dataset was created from the same base PPI-graph used to create the PPI-adjacency dataset, but now using as features only 18 graph measures applied to the 850 ageing-related genes within the PPI graph. These measures are classified as centrality- and non-centrality-based and were computed using the *R*’s igraph and *Centricerve* libraries (Csardi and Nepusz, 2006; Jalili et al., 2015, 2016). Following these libraries’ documentation, the centrality measures used in this work were: Leverage, Markov, Maximum neighborhood component, Closeness, Betweenness, Laplacian, Diffusion, Semilocal, Subgraph, Geokpat, Eigenvalue, Eccentricity, Degree and Lobb centralities. In addition, we used the Kcore, CR-ratio, Clustering Coefficient and Topological Coefficient as non-centrality-based measures. A detailed description of all these measures, except CR-ratio, is available on the *Centiserve* webpage (Jalili et al., 2015). The CR-ratio is the ratio of the number of CR-related direct neighbours of the queried gene over the total number of neighbours of the queried gene (*i.e*., it describes what percentage of the queried-gene’s direct neighbours are CR-related). Figure 2.C represents this dataset by displaying a graph whose nodes are coloured based on their degree centrality (the larger the number of neighbours, the higher the degree centrality and the darker the nodes’ colour).

#### 2.1.7 GO terms Dataset

In this dataset, each binary feature represents a specific GO term with which each ageing-related gene may or may not be associated. To accomplish this, a *BioMart* query retrieved a list of the Biological Process GO terms associated with the ageing-related genes, yielding 8640 different GO terms. The retrieved GO terms form a hierarchical structure with two properties: 1) if a gene is associated to a specific GO term, it will also be associated with all its ancestors (*i.e*., more general GO terms); and 2) if a gene is not associated to a given GO term, then the gene is not associated to any of its descendants. The GO term GO:0008150 (named ‘Biological Process’) is the root of the hierarchy, *i.e*., it is an ancestor of all other GO terms in the dataset. The *R*’s *GO.db* package [Carlson, 2019] was used to retrieve all the ancestors of the originally retrieved GO terms. Those ancestors were then merged to this dataset as new predictive features. This process increased the number of GO terms across all the ageing-related genes from 4877 (*i.e*., without ancestors) to 8640 (*i.e*., containing both the original retrieved GO terms and their ancestors). Each GO term (considering both originally retrieved and ancestor GO terms) is represented as a binary feature, indicating whether or not a gene (instance) is annotated with that GO term. Out of the 1137 ageing-related genes, 13 genes were not associated with any GO term in *BioMart* and were removed. This produced a dataset composed of 1124 ageing-related genes: 114 labelled as *Ageing*_*CR*_ and 1010 labelled as *Ageing*_*NotCR*_. Finally, due to the hierarchical structure of GO terms, any gene associated with a fixed GO term is also associated with all of the term’s ancestors. Figure. 2.D illustrates this phenomenon by indicating that association with a fixed GO term (red node) implies association with all of its ancestors (orange nodes).

#### 2.1.8 GTEx Dataset

The median expression levels of human genes across 55 different anatomical tissues were retrieved from the GTEx database (Carithers et al., 2015) (Analysis V8 database) by downloading the file corresponding to the median gene-level Transcripts Per Million (TPM) by tissue. Then, only ageing-related genes were retained, resulting in 1111 ageing-related genes: 114 labelled as *Ageing*_*CR*_-related and 997 labelled as *Ageing*_*NotCR*_-related. The tissues’ median TPM scores were then used as predictive features. A graphical representation of this dataset is illustrated in Figure 2.E through a heatmap of the mean expression of each single gene across different tissues.

#### 2.1.9 Co-expression Dataset

The *GeneFriends* database (Dam et al., 2015) was used to generate co-expression profiles for 1048 ageing-related genes across a set of 44,946 genes that included both the 1048 ageing-related and other genes. The goal was to determine whether the co-expression profile of key genes across the ageing-related genes contributes to the association of certain age-related genes with CR. This dataset contained 106 and 942 *Ageing*_*CR*_- and *Ageing*_*NotCR*_-related genes, respectively. Figure 2.F illustrates a representation of this dataset, where the correlation between the expression of two genes across different samples is depicted.

#### 2.1.10 Protein-Descriptor Dataset

This dataset contains numeric features associated with the proteins encoded by the ageing-related genes. Since each gene may code for either one or more proteins, this dataset differs from others in the sense that it provides information on ageing-related proteins rather than genes. The names and sequences of human proteins were obtained using the *proteins* command in *R*’s *ensembldb* library, with the database *EnsDb.Hsapiens.v86* (Rainer, 2017; Rainer et al., 2019) as the source. 1109 ageing-related genes encoded 6180 ageing-related proteins, from where 115 genes and 514 proteins were designated as *Ageing*_*CR*_-related, while the remaining 994 genes and 5666 proteins as *Ageing*_*NotCR*_-related. Additionally, features were computed from the amino acid sequences of the *proteins* using the *R*’s *protr* and Peptides packages (Xiao et al., 2015; Osorio et al., 2015), which resulted in the features discussed in Supplementary materials (S.1.2).

#### 2.1.11 Whole-Datasets

Two datasets were created that combine features from PathDIP, KEGG-Pertinence, KEGG-Influence, PPI adjacency matrix, PPI graph measures, GO terms, GTEx expression data, and Co-expression datasets. The protein-descriptors dataset was not included as it only provides information on proteins rather than genes, which complicates the gene-based merging process as proteins do not always have a one-to-one relationship with genes. We coined the term ‘WholeDataset’ to refer to the resulting dataset that combines all of the aforementioned features, yielding a total of 63,099 features and 1,137 ageing-related genes: 115 labelled as *Ageing*_*CR*_ and 1022 labelled as *Ageing*_*NotCR*_.

Since the merged datasets had different numbers of ageing-related genes, the WholeDataset contained data gaps for ageing-related genes whose features were not annotated across all the datasets. We addressed this issue using two approaches, namely, imputation and intersection, which are described next and illustrated in Figure 2.G, where the combination of datasets results in genes with missing data (purple cells), which are then imputed (green cells) or removed to leave only genes containing features from all the datasets (intersected genes).

##### Whole-Dataset-imputation

In this approach we used a 5 Nearest Neighbors (5NN) imputation method. For each ageing-related gene G that is missing a value for feature F, this method first determines the top five ageing-related genes that have a known value for F and are most similar (have the smallest Euclidean distance) to G in the training set. The Euclidean distance is computed using all the features for which the values of G are known. If F is a continuous feature, its missing value in G is imputed using the mean value of F across the 5NN of G in the training set. If F is binary, the missing value in G is imputed using the mode of F (*i.e*., its most frequent value) across the 5NN of G in the training set.

##### Whole-Dataset-intersection

In this approach we only retained the ageing-related genes that are present across every single dataset, except the protein-descriptors dataset. This guarantees the absence of any missing feature values. However, information is lost since only about half of the ageing-related genes had known values for all the features. The resulting dataset contained 628 ageing-related genes: 72 labelled as *Ageing*_*CR*_-related and 556 labelled as *Ageing*_*NotCR*_-related.

### 2.2 Machine Learning

This work focuses on decision tree-based ensembles, which are a type of powerful ML technique that combine the predictions of several base learners (decision trees) in order to improve predictive accuracy over a single base learner. This type of ensembles is usually categorised into two broad groups: (1) Bagging methods, where each base learner is trained independently from the others – so, the base learners are conceptually trained in parallel. In bagging methods, the predictive accuracy is usually improved due to the reduction of the variance in the ensemble’s predictions, by comparison with the variance in the predictions of a single base learner. (2) Boosting methods, where the base learners are trained sequentially, and each base learner is trained with instance weights that are determined in order to correct the errors of previous base learners in the sequence. This tends to reduce the bias in the predictions (Pedregosa et al., 2011).

One challenge in this work is that *Ageing*_*NotCR*_-related genes are roughly tenfold more numerous than *Ageing*_*CR*_-related genes, resulting in an imbalanced data that biases ML predictions towards *Ageing*_*NotCR*_-related genes. To address this, under-sampling of the majority class (*Ageing*_*NotCR*_-related genes) was performed for each of their base learners’ training set. Hence, after under-sampling each training set (for each base learner) has the same number of *Ageing*_*CR*_-related and *Ageing*_*NotCR*_-related genes. Two of the ML algorithms we used, Balanced Random Forests (BRF) and Easy Ensemble Classifier (EEC), are bagging and boosting methods, respectively. Both BRF and EEC were implemented using the *Python* package *Imbalanced-learn* (Lemaître et al., *2017)*. Additionally, XGBoost (XGB) (Chen and Guestrin, 2016) and CatBoost (CAT) (Dorogush et al., 2018) were used, as they are both high-performance opensource libraries for gradient boosting in decision trees. Figure 3.A illustrates these algorithms graphically. The four *techniques were* run with *random _state* set to 42. Further details on data preprocessing are available in Supplementary materials (S.2.1).

**Figure 3:**
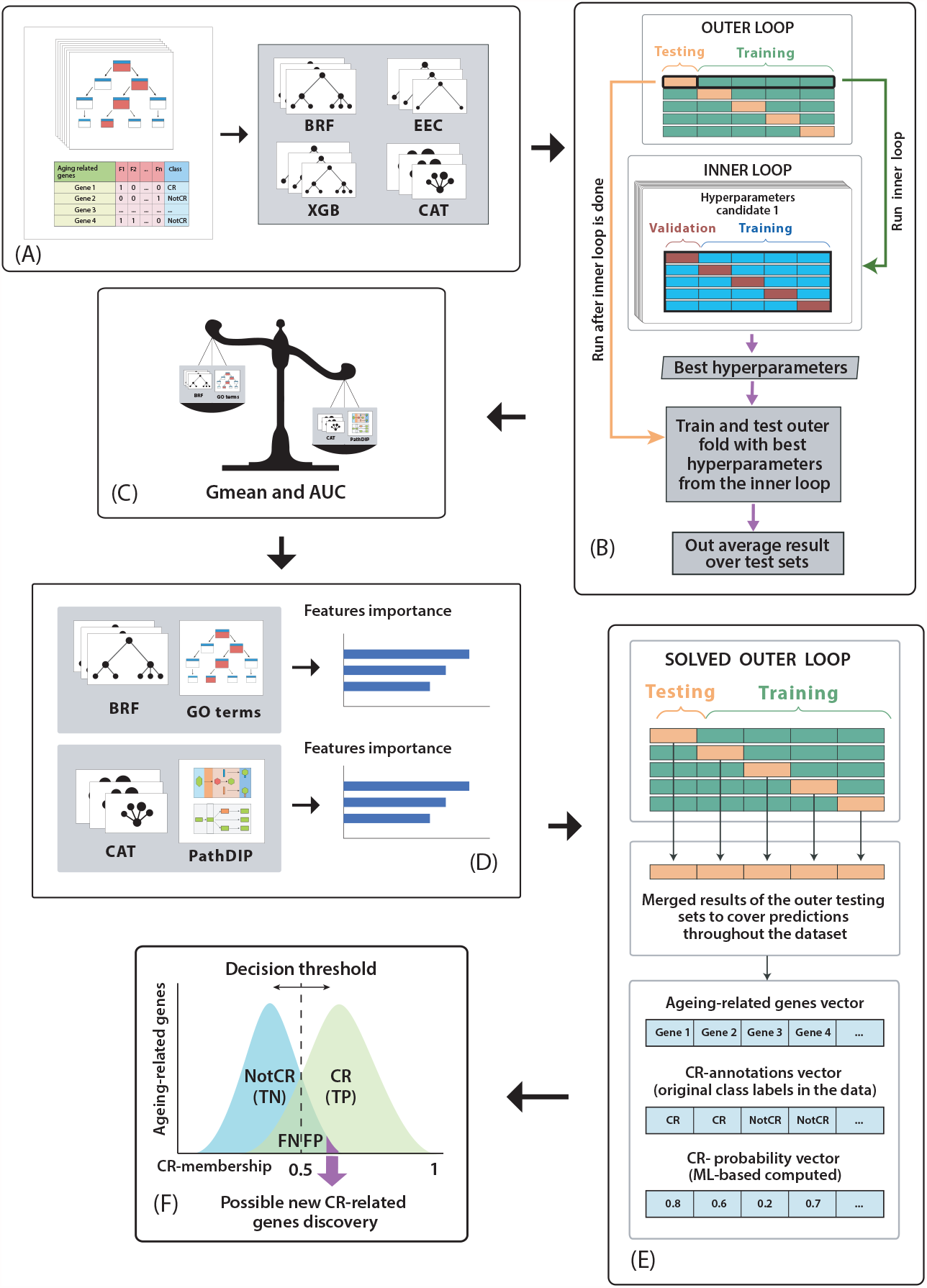
Summary of ML methods. **A**. Selection of each possible {Dataset, ML algorithm} model. **B**. Evaluation of each model through the nested-CV. Only 5 outer folds are depicted to facilitate visualization. **C**. Comparison of the predictive power of all the {Dataset, ML algorithm} models using Gmean and AUC. **D**. Computation of feature importance analysis in the two best performing models from different datasets. **E**. Construction of the *CR-annotations* and the *CR-probability* vectors. **F**. Possible CR-related inference from *Ageing*_*NotCR*_-related genes strongly predicted as CR-related. TN, TP, FN and FP refer to True Negatives, True Positives, False Negatives and False Positives, respectively.

#### 2.2.1 Predictive accuracy calculation

Predictive accuracy was calculated by using a nested cross-validation (CV) procedure (a common approach in ML), as follows. To implement the outer 10-fold cross-validation, the dataset instances (ageing-related genes) are randomly divided into 10 outer folds of roughly the same size. Then each ML algorithm is run 10 times, each using a different outer fold as the testing set and all the other 9 outer folds as the training set. Before each run of an algorithm, however, its hyperparameters are tuned by an inner 5-fold cross-validation. To implement this, each training set of the outer CV is randomly divided into 5 inner folds of roughly the same size. Then, for each candidate configuration of hyperparameter settings of the algorithm, the algorithm is run 5 times, each time using a different inner fold as the validation set (to measure predictive accuracy) and the other 4 inner folds as a reduced training set. The predictive accuracy of each candidate algorithm configuration is computed as the average accuracy over the 5 validations sets, and then the algorithm configuration with the highest average predictive accuracy is chosen as the best configuration for the current iteration of the outer CV. Next, the algorithm with that best configuration is applied to the full training set of the current iteration of the outer CV, and the learned classifier is evaluated on the current testing set. Finally, this whole process is repeated for the 10 iterations of the outer CV, and the predictive accuracy of the algorithm is computed as the average of the 10 values of predictive accuracy over the 10 testing sets of the outer CV. Note that hyperparameter optimization is performed by the inner CV using only the training set (*i.e*., not using the testing set), which is always reserved for measuring generalisation performance. A graphical representation of the nested CV procedure is depicted in in Figure 3.B. (which only displays five outer folds for ease of visualization). In this work, the inner loop was implemented using the *scikit-learn*’s *GridSearchCV* command (Pedregosa et al., 2011), with *random_state* = 42 and hyperparameters as stated in Supplementary materials (S.2.2).

The performance metric used for hyperparameter tuning during the inner-loop of the nested CV was the Geometric Mean (Gmean), defined in equation (1):

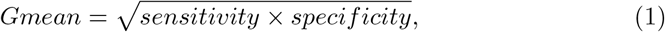

where sensitivity is the percentage of *Ageing*_*CR*_-related genes (*i.e*., the minority class) that were correctly predicted as *Ageing*_*CR*_-related, and specificity is the percentage of *Ageing*_*NotCR*_-related genes (*i.e*., the majority class) that were correctly predicted as *Ageing*_*NotCR*_-related. Since sensitivity and specificity take values in the [0, 1] interval, so does Gmean. Gmean is suited for class-unbalanced problems as this metric measures the balance between classification performances on both the majority and minority classes (Ri and Kim, 2020). The closer Gmean is to 1, the better is the classification.

The predictive performance of the final models on the testing sets of the outer CV was evaluated by two measures (Figure 3.C): (a) Gmean of sensitivity and specificity, and (b) Area Under the Receiver Operating Characteristic Curve (AUC), which is an overall summary of predictive accuracy. AUC also takes values in [0, 1], where 1 is the ideal value (indicating that all predictions were correct), and an AUC value of 0.5 corresponds to random predictions.

A special form of nested CV procedure was applied to the protein-descriptors dataset, as follows. The sequences of ageing-related proteins encoded by a single ageing-related gene are highly correlated. As a result, the testing and training sets of the proteins dataset are also likely to be highly correlated, impeding a fair measure of predictive accuracy. To address this issue, the inner and outer folds of the nested-CV were performed at the gene level rather than at the protein level. This was accomplished by directly applying the nested-CV splitting to all the ageing-related genes containing proteins in the *EnsDb.Hsapiens.v86* database (Rainer, 2017; Rainer et al., 2019). Following that, the corresponding proteins for each of these ageing-related genes were retrieved. Proteins encoded by the genes in the outer training, outer testing, inner training and inner validation sets were then used to create their corresponding data subsets of the protein-descriptors dataset.

#### 2.2.2 Feature Importance Calculation

We calculated the feature importance for the best learned models (Figure 3.D). To accomplish this, we used 100% of the ageing-related genes (instances) of the dataset under study as the training set, which ensured that the features importance were calculated using all of the data available, maximising the quality of the feature importance calculation. No instances were withheld for testing purposes, as this task’s objective was not to determine predictive accuracy (already determined by the nested-CV procedure) but to compute the features’ predictive relevance.

The importance of BRF’s features is determined by using the default Gini index of class impurity, which calculates how well a split separates the samples of the two classes in each node of a decision tree. A feature’s importance is basically given by the weighted average of the reduction of the Gini index across the tree nodes labelled with that feature, with weights proportional to the number of instances split by that feature (Menze et al., 2009). On the other hand, EEC, XGB, and CAT calculate a feature’s relevance through the permutation method, which compares the model’s predictive accuracy on the original data vs the model’s accuracy on a dataset with a random permutation of that feature’s values, so that the extent of the drop in the model’s accuracy after the random permutation indicates how much the model is dependent on the feature. Finally, in order to compare the results of the two feature importance methods, the resulting feature rankings were scaled from their original values to the [0, 100] interval, where 100 represents the most important feature and 0 represents no relevance.

#### 2.2.3 New CR-related genes inference

Each learned classification model outputs a probability that a gene belongs to the *Ageing*_*CR*_-related class. For converting a predicted probability into a class label we use a classification threshold of 0.5, *i.e*., any gene with a predicted *Ageing*_*CR*_-related probability less than 0.5 is predicted to be *Ageing*_*NotCR*_-related, whereas any gene with a predicted probability greater than 0.5 is predicted to be *Ageing*_*CR*_-related. Although CR-related predictions are binarized by the threshold, the CR-related probabilities remain informative as a measure of the prediction’s certainty, from the model’s viewpoint. For instance, if two given genes, *A* and *B*, have CR-related probabilities of 0.6 and 0.9, respectively, both are classified as CR-related; but from the model’s perspective, gene *B* is more reliably related with CR.

Hence, it is possible to infer novel *Ageing*_*CR*_-related genes by identifying, among all the genes annotated in the dataset as *Ageing*_*NotCR*_-related genes, which ones have the greatest predicted *Ageing*_*CR*_-related probabilities, which are the strongest false positives (FP) genes. After all, the *Ageing*_*NotCR*_-related class label annotation in the dataset is not very reliable in general, because it basically means that there is no evidence for *Ageing*_*CR*_-relatedness in the literature, and absence of evidence is not the same as evidence for absence of *Ageing*_*CR*_-relatedness. Hence, the strongest FP genes are good candidate targets for future experiments to determine *Ageing*_*CR*_-related genes.

Keeping in mind that the nested-CV’s outer testing sets do not overlap and that they collectively include all the genes (instances) in the dataset, we created, for each of the best performing Dataset, ML algorithm combinations (*i.e*., the best models), two vectors: a *CR-probability vector* combining the predicted CR-related probabilities from all the outer testing splits, and a *CR-annotations vector* by combining the original annotations of class labels in the data from all the outer splits (Figure 3.E). Next, based on these two vectors, we retained only *Ageing*_*NotCR*_-related genes with CR-probability equal to or greater than 0.5 (*i.e*., FP genes), meaning that they were classified as CR-related by the model but lack a CR-related annotation in the dataset (based on the literature). We identified the top 10 FP genes with greater CR-probability for each of the best performing models and discussed their potential as candidate CR-related genes for confirmation in further wet-lab experiments. (Figure 3.F).

Finally, we looked for common top CR-related genes candidates among the shared ageing-related genes between the two strongest models, namely {GO terms, BRF} and {PathDIP, CAT}, as described in the Results section. To do so, we originally computed a common-ranking by averaging the CR-probability scores of common genes in both models and then sorted the genes in descendent order based on the averaged score. This approach had, however, one issue as the CR-probability density distributions of both strongest models lied in different intervals ([0.35,0.75] in {GO terms, BRF} while [0,1] in {PathDIP, CAT}, as described in Results). Consequently, each computed average was biased towards the distribution with the most extreme values (*i.e*., genes with the greatest/lowest CR-probability scores in {PathDIP, CAT} will have stronger influence when averaging than genes with greatest/lowest CR-membership score in {GO terms, BRF}). With the aim to provide a similar comparison scale for the top CR-related candidates in both models while considering the distribution shape, we mapped the CR-membership scores of all genes in both {GO terms, BRF} and PathDIP, CAT models to the [0,1] interval and then retrieved common genes in both models to compute CR-probability arithmetic averages, highlighting *Ageing*_*NotCR*_-related genes with the highest probabilities of being *Ageing*_*CR*_-related genes from both best models’ perspectives.

### 2.3 Statistical analysis

To report on the most important features, two statistical analysis tests were used (with a significance level of 0.01): a Two-Proportions Z-Test for binary features and a T-test for continuous features. The use of the test for binary features is based on the concept of a feature’s positive value, which is defined as the presence of annotation (*e.g*., a GO term annotation) for a gene, as opposed to the absence of that annotation (the negative value of the feature).

The test for binary features was designed to determine whether the proportion of *Ageing*_*CR*_-related genes associated with a particular relevant feature’s positive value (*i.e*., the ratio of *Ageing*_*CR*_-related genes associated with the relevant feature’s positive value over the total number of *Ageing*_*CR*_-related genes in the dataset) is significantly different than the proportion of *Ageing*_*NotCR*_-related genes associated with the same relevant feature’s positive value. For continuous features, the test determined whether the mean value of the feature across all *Ageing*_*CR*_-related genes was significantly different than the mean value of the feature across all *Ageing*_*NotCR*_-related genes.

## 3 Results

This section first highlights the best performing combinations of ML algorithms and datasets. Top relevant features are then presented. Finally, the predicted top CR-associated genes candidates are reported.

### 3.1 Predictive Accuracy Results

The AUC and Gmean values obtained by BRF, EEC, XGB, CAT are compared in Table 1. For each dataset (feature type) and predictive accuracy measure, the best result from the four algorithms is highlighted in boldface. GTEx and co-expression were the least predictive feature types, yielding results comparable to random predictions (AUC close to 0.5), whereas GO terms and PathDIP were the most predictive: GO terms had the highest average AUC (0.83) and the second highest Gmean (0.75) across the four algorithms; whilst PathDIP had the highest average Gmean (0.76) and the second highest average AUC (0.81). BRF got the highest AUC values overall, whereas the highest Gmean values were more distributed across all algorithms, with higher means for EEC and CAT.

**Table 1:** Gmean and AUC scores for the BRF, EEC, XGB, and CAT algorithms across all the 11 datasets (feature types). The average Gmean and AUC of each algorithm across all the datasets is shown in the last row; and the average Gmean and AUC of each feature type across all four algorithms is shown in the last two columns.

| Feature Type | Num. of features | Num. of instances | Gmean |  |  |  | AUC |  |  |  | Mean |  |
| --- | --- | --- | --- | --- | --- | --- | --- | --- | --- | --- | --- | --- |
|  |  |  | BRF | EEC | XGB | CAT | BRF | EEC | XGB | CAT | Gmean | AUC |
| PathDIP | 1640 | 986 genes | 0.76 | 0.75 | 0.75 | **0.77** | 0.81 | 0.8 | 0.8 | **0.83** | **0.76** | 0.81 |
| KEGG-Pertinence | 312 | 799 genes | 0.73 | 0.7 | **0.75** | 0.72 | **0.8** | 0.71 | 0.76 | 0.73 | 0.73 | 0.75 |
| KEGG-Influence | 1770 | 799 genes | **0.7** | **0.7** | 0.67 | 0.67 | **0.78** | 0.71 | 0.69 | 0.68 | 0.69 | 0.72 |
| PPI-measures | 18 | 850 genes | 0.64 | 0.59 | **0.65** | **0.65** | **0.69** | 0.6 | 0.65 | 0.67 | 0.63 | 0.65 |
| PPI-adjacency | 5718 | 850 genes | 0.62 | 0.61 | 0.59 | **0.65** | **0.72** | 0.63 | 0.64 | 0.66 | 0.62 | 0.66 |
| GO terms | 8640 | 1124 genes | **0.76** | 0.75 | 0.74 | 0.74 | **0.84** | 0.83 | 0.83 | 0.81 | 0.75 | **0.83** |
| GTE<sub>x</sub> | 55 | 1111 genes | 0.5 | **0.52** | 0.5 | 0.51 | 0.5 | **0.53** | 0.52 | 0.52 | 0.51 | 0.52 |
| Co-expression | 44,946 | 1048 genes | 0.5 | **0.57** | 0.42 | 0.56 | 0.52 | **0.57** | 0.54 | 0.54 | 0.51 | 0.54 |
| Proteins-Descriptors | 156 | 6180 proteins (from 1109 genes) | 0.45 | 0.61 | **0.62** | 0.6 | 0.65 | **0.67** | **0.67** | 0.63 | 0.57 | 0.66 |
| Whole-Dataset Imputation | 63,099 | 1137 genes | **0.67** | 0.65 | 0.6 | 0.66 | **0.72** | 0.66 | 0.65 | 0.67 | 0.65 | 0.68 |
| Whole-Dataset Intersection | 63,099 | 628 genes | 0.61 | **0.66** | 0.56 | 0.64 | 0.64 | **0.67** | 0.63 | 0.66 | 0.62 | 0.65 |
| Mean | - | - | 0.63 | **0.65** | 0.62 | **0.65** | **0.70** | 0.67 | 0.67 | 0.67 | 0.64 | 0.68 |

By defining a model as a combination of a dataset and the classification algorithm that runs over it Dataset, ML algorithm, the model with the highest Gmean (0.77) was {PathDIP, CAT}, closely followed by {GO Terms, BRF} and {PathDIP, BRF}, both with a Gmean of 0.76. Regarding AUC results, the best model was {GO Terms, BRF} with an AUC of 0.84, closely followed by {GO terms, EEC} and {PathDIP, CAT}, both with an AUC of 0.83. Since {PathDIP, CAT} and {GO Terms, BRF} achieved complementary and notably close first and second places regarding Gmean and AUC, so they are both the most predictive models overall.

Table 2 reports sensitivity and specificity results, as measures of predictive accuracy for *Ageing*_*CR*_-related genes and *Ageing*_*NotCR*_-related genes, respectively. The highest mean sensitivity and specificity values across all four algorithms were obtained by PathDIP and Proteins-Descriptors, respectively, as shown in the last two columns of the table. There was no strong winner algorithm in terms of sensitivity, but XGB obtained by far the worst sensitivity values overall, as shown in the last row of the table. On the other hand, XGB achieved in general the highest specificity values, implying more accurate predictions for *Ageing*_*NotCR*_-related genes, the majority class. ROC curves of all the ML algorithms in the two strongest datasets, as well as confusion matrices of the two strongest models, {GO Terms, BRF} and {PathDIP, CAT}, are depicted in supplementary materials (Figures S1 and S2, respectively).

**Table 2:** Sensitivity (*Ageing*_*CR*_ prediction quality) and specificity (*Ageing*_*NotCR*_ prediction quality) values for the BRF, EEC, XGB, and CAT algorithms across all the datasets (feature types). The average sensitivity and specificity of each algorithm across all the datasets is shown in the last row; and the average Gmean and AUC of each feature type across all four algorithms is shown in the last two columns.

| Feature Type | Num. of features | Num. of instances | Sensitivity |  |  |  | Specificity |  |  |  | Mean |  |
| --- | --- | --- | --- | --- | --- | --- | --- | --- | --- | --- | --- | --- |
|  |  |  | BRF | EEC | XGB | CAT | BRF | EEC | XGB | CAT | Sens. | Spec. |
| PathDIP | 1640 | 986 genes | **0.87** | 0.77 | 0.58 | 0.77 | 0.67 | 0.7 | **0.93** | 0.77 | **0.75** | 0.77 |
| KEGG-Pertinence | 312 | 799 genes | 0.74 | 0.70 | 0.75 | 0.72 | 0.7 | 0.71 | **0.76** | 0.73 | 0.73 | 0.73 |
| KEGG-Influence | 1770 | 799 genes | **0.75** | 0.70 | 0.67 | 0.67 | 0.66 | **0.71** | 0.69 | 0.68 | 0.70 | 0.69 |
| PPI-measures | 18 | 850 genes | **0.64** | 0.62 | 0.6 | 0.57 | 0.65 | 0.57 | 0.71 | **0.77** | 0.61 | 0.68 |
| PPI-adjacency | 5718 | 850 genes | **0.65** | 0.52 | 0.5 | **0.65** | 0.62 | 0.74 | **0.78** | 0.67 | 0.58 | 0.70 |
| GO terms | 8640 | 1124 genes | **0.85** | 0.80 | 0.48 | 0.69 | 0.67 | 0.72 | **0.94** | 0.78 | 0.71 | 0.78 |
| GTE<sub>x</sub> | 55 | 1111 genes | 0.54 | 0.55 | 0.45 | **0.57** | 0.48 | 0.51 | **0.57** | 0.48 | 0.53 | 0.51 |
| Co-expression | 44,946 | 1048 genes | 0.58 | **0.61** | 0.21 | 0.56 | 0.45 | **0.86** | 0.85 | 0.54 | 0.49 | 0.68 |
| Proteins-Descriptors | 156 | 6180 proteins (from 1109 genes) | 0.25 | 0.43 | 0.45 | **0.48** | 0.86 | **0.91** | 0.88 | 0.77 | 0.40 | **0.86** |
| Whole-Dataset Imputation | 63,099 | 1137 genes | 0.69 | 0.68 | 0.43 | **0.71** | 0.65 | 0.63 | **0.87** | 0.64 | 0.63 | 0.70 |
| Whole-Dataset Intersection | 63,099 | 628 genes | 0.60 | 0.72 | 0.39 | **0.75** | 0.63 | 0.62 | **0.87** | 0.56 | 0.62 | 0.67 |
| Mean | - | - | **0.65** | **0.65** | 0.50 | **0.65** | 0.64 | 0.70 | **0.80** | 0.67 | 0.61 | 0.70 |

### 3.2 Feature Importance Results

The top-5 most relevant features in each of the two most predictive models, {GO terms, BRF} and {PathDIP, CAT}, are shown in Tables 3 and 4, respectively. The column *Score* in these tables indicates the relative importance of the features, in the range from 0 (no relevance) to 100 (maximal relevance).

**Table 3:** Top-five most predictive GO terms in the {GO terms, BRF} model.

| Features Importance for the {GO-Terms, BRF} model |  |  |  |  |  |
| --- | --- | --- | --- | --- | --- |
| Feature | Definition | Score | $Ageing_{CR}$ (%) | $Ageing_{NotCR}$ (%) | p-value |
| GO:0055114 | oxidation-reduction process | 100 | 19.3%<br>{22/114} | 11.58%<br>{117/1010} | 0.013 |
| GO:0007188 | adenylate cyclase-modulating G protein-coupled receptor signalling pathway | 70.965 | 13.16%<br>{15/114} | 0.3%<br>{3/1010} | **9.72e-24** |
| GO:1904659 | glucose transmembrane transport | 67.831 | 10.53%<br>{12/114} | 0%<br>{0/1010} | **2.40e-23** |
| GO:0007186 | G protein-coupled receptor signalling pathway | 53.516 | 15.79%<br>{18/114} | 2.77%<br>{28/1010} | **7.73e-11** |
| GO:0008643 | carbohydrate transport | 51.516 | 9.65%<br>{11/114} | 0.1%<br>{1/1010} | **2.24e-19** |

**Table 4:**
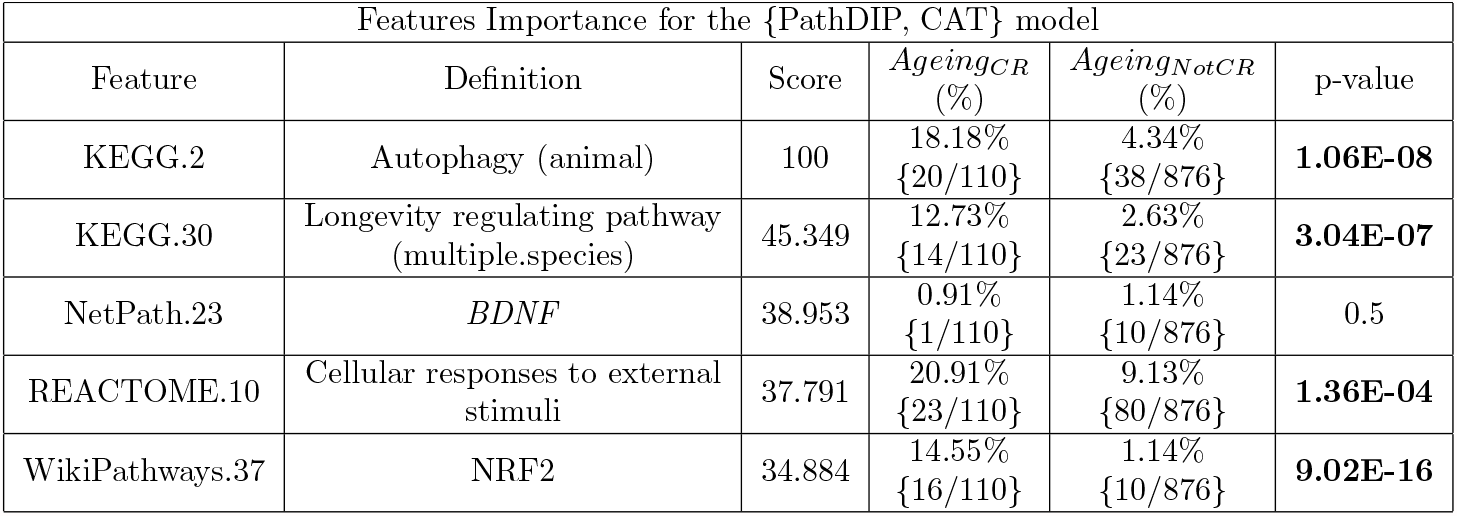
Top-five most predictive pathways in the {PathDIP, CAT} model

The columns *Ageing*_*CR*_ and *Ageing*_*NotCR*_ denote the percentage of *Ageing*_*CR*_- and *Ageing*_*NotCR*_-related genes with the GO term or pathway in the corresponding row (*i.e*., the percentage of genes with the feature’s positive value). In addition, a proportion is provided in brackets where the numerator indicates the number of *Ageing*_*CR*_- or *Ageing*_*NotCR*_-related genes with the positive feature value, while the denominator indicates the total number of *Ageing*_*CR*_- or *Ageing*_*NotCR*_-related genes, in the dataset.

The *p-value* columns in Tables 3 and 4 indicate the results of the statistical tests applied to detect whether the values in the *Ageing*_*CR*_ column are significantly different from those in the *Ageing*_*NotCR*_ column as explained in Section 2.3. Significant p-values are denoted by bold text.

Interestingly, among the GO terms in Table 3, the most relevant one, oxidation-reduction process, was the only one that failed to achieve a significant difference in terms of the proportion of *Ageing*_*CR*_- and *Ageing*_*NotCR*_-related genes. Nonetheless, this term is worth highlighting due to it having the highest proportion of occurrence (19.3%) in *Ageing*_*CR*_-related genes among all 5 GO terms in this table. The remaining GO terms, which are related to GPCRs and carbohydrate transport, clearly have a stronger association with *Ageing*_*CR*_-related genes, as each of them occurred in about 10%–16% of the *Ageing*_*CR*_-related genes while occurring in less than 3% of the *Ageing*_*NotCR*_-related genes.

Table 4 reports the top PathDIP features. Note that the score for the second best pathway, longevity regulating pathway, is much lower than the score for the best pathway, autophagy. The only feature with no significant difference in its percentage of occurrence in *Ageing*_*CR*_- and *Ageing*_*NotCR*_-related genes was brain-derived neurotrophic factor (*BDNF*). The cellular responses to external stimuli pathway contained the greatest proportion of occurrence (20.9%) in *Ageing*_*CR*_-genes.

### 3.3 CR-associated gene prediction

The two most predictive models, {GO terms, BRF} and {PathDIP, CAT}, were learned from datasets with 1124 and 986 ageing-related genes, respectively. Figures 4.A and 4.B show the distribution of *Ageing*_*CR*_- and *Ageing*_*NotCR*_-related genes across different CR-probability values for {GO terms, BRF} and {PathDIP, CAT}, respectively. The CR-probability distributions in {GO terms, BRF} span a much narrower window, near [0.35, 0.70], than distributions in {PathDIP, CAT}, near [0, 1]. Additionally, both models’ *Ageing*_*NotCR*_-related genes share a maximal density point that is relatively close to the 0.5 threshold, yet on the *Ageing*_*NotCR*_-predicted side (0.45 points). This point is notably denser (about 6 folds) and narrower in {GO terms, BRF} than it is in {PathDIP, CAT}.

**Figure 4:**
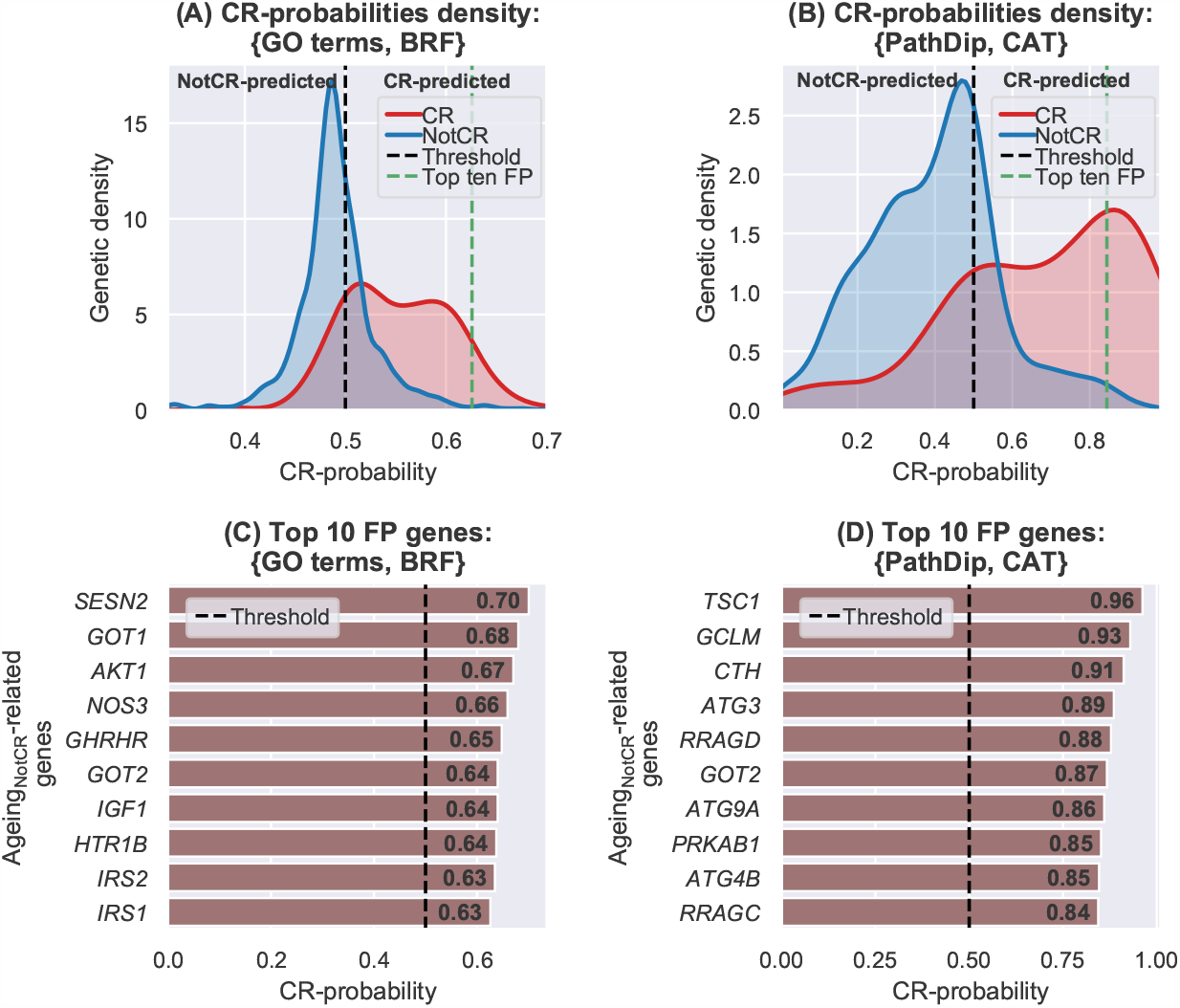
**A** and **B**, CR-probabilities density distribution of {GO terms, BRF} and {PathDIP, CAT}, respectively. The green dashed line represents the beginning of the top 10 False Positives (FP), which are identified as candidate CR-associated genes. **C** and **D**, top 10 FP genes in {GO terms, BRF} and {PathDIP, CAT}, respectively. The classification threshold (0.5) is represented by the dashed black line.

Regarding the top CR candidate genes, further analysis of Figures 4.A and 4.B, indicates that, for both models, the top CR-probabilities across *Ageing*_*NotCR*_-related genes achieved similar scores that top CR-probabilities in *Ageing*_*CR*_-related genes, but are much less numerous. Figures 4.C and 4.D depict the top ten CR-candidate genes in the {GO terms, BRF} and {PathDIP, CAT} models, respectively. Even if some of the top genes predicted by both models may have a proclivity for not detecting unknown CR-relationship, it is remarkable that among their top ten CR-candidate genes, only *GOT2* overlapped.

Hypothesising that pertinence to the common set of top-10 CR-candidate genes in both models increases likelihood of accurate CR-relatedness prediction, we performed a joint analysis of the top -10 CR-candidate genes (Methods 2.2.3). Briefly, we normalised the range of both models’ CR-probability distributions and then retained common ageing-related genes (976 genes, 872 of which are *Ageing*_*NotCR*_-related) in order to compute, for each common ageing-related gene, an arithmetic average of both normalised CR-probabilities. Then, we sort genes under a criteria that considers both a similar CR-membership range for the two models and their distribution shapes. The correlation between the normalised CR-probabilities assigned by the {GO terms, BRF} and {PathDIP, CAT} models was only moderated, being smaller across *Ageing*_*NotCR*_-related genes, increasing throughout *Ageing*_*CR*_-related genes and yielding the highest score when using all ageing-related genes (Figure 5).

**Figure 5:**
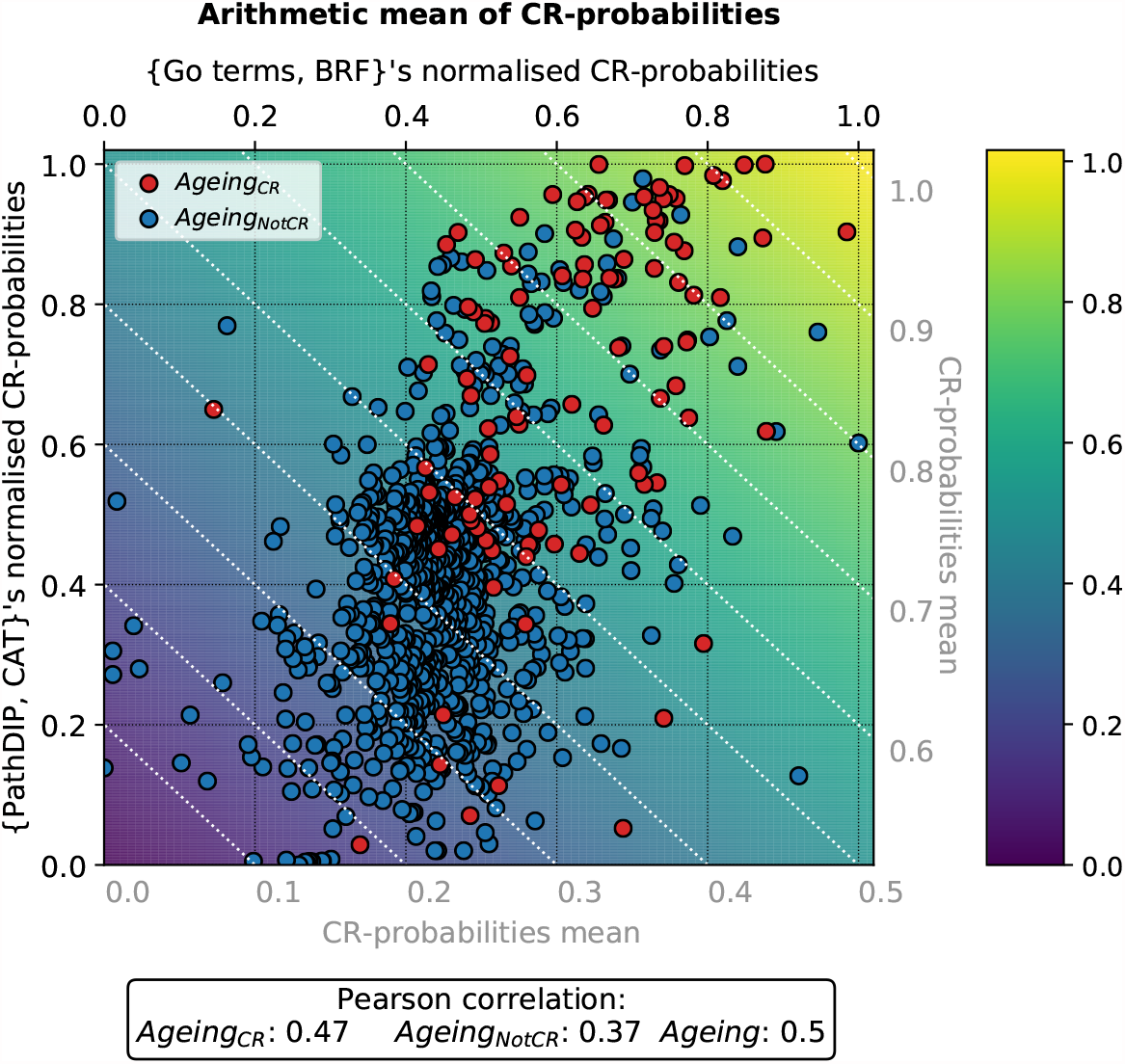
Correlation of normalised CR-probabilities assigned by {GO terms, BRF} and {PathDIP, CAT} models. The arithmetic mean of both models’ normalised CR-probabilities is depicted through the background color while the white dotted lines represent points in the plane with fixed mean values indicated by the grey ticks. The Pearson correlation score between the the normalised CR-probabilities applied by the two models is depicted in the text box at the bottom of the figure.

Table 5 depicts both models’ common *Ageing*_*NotCR*_-related genes whose averaged normalised CR-probability exceed 0.8. Note that all of these genes, namely, *GOT2, GOT1, TSC1, CTH, GCLM, IRS2*, and *SENS2* ; appeared in the top-10 CR-related candidates of at least one of the two most predictive models. The top gene in 5, Glutamic-Oxaloacetic Transaminase 2 (*GOT2*), appeared within the set of top six ranking genes of both models, indicating that its possible CR-relatedness could be similarly inferred from either biological pathways or biological processes GO terms. Insulin Receptor Substrate 2 (*IRS2*) and especially Glutamic-Oxaloacetic Transaminase 1 (*GOT1*) got high CR-probabilities in the {GO terms, BRF} model, and a moderately high probability in {PathDIP, CAT}. Moreover, Sestrin 2 (*SENS2*), the gene with the highest CR-probability in the {GO terms, BRF} model, also reached the list but it was, by far, the gene with lowest CR-probability in {PathDIP, CAT}, among all genes in Table 5. Finally, TSC Complex Subunit 1 (*TSC1*), glutamate-cysteine ligase regulatory subunit (*GCLM*) and Cystathionine Gamma-Lyase (*CTH*) got the three highest CR-probability scores in {PathDIP, CAT}, and also relatively high probabilities in {GO terms, BRF}.

**Table 5:** Top CR-related gene candidates jointly defined by the two strongest models. Top-association ranking is provided relative to the number of common *Ageing*_*NotCR*_-related genes.

| Joint Rank | Gene | Normalised CR-probability |  |  | Ranking |  |
| --- | --- | --- | --- | --- | --- | --- |
|  |  | Mean. Score | {GO terms, BRF} | {PathDIP, CAT} | {GO terms, BRF} | {PathDIP, CAT} |
| 1 | *GOT2* | 0.861 | 0.840 | 0.882 | 5th | 6th |
| 2 | *GOT1* | 0.853 | 0.946 | 0.760 | 2th | 35th |
| 3 | *TSC1* | 0.847 | 0.714 | 0.979 | 19th | 1th |
| 4 | *CTH* | 0.846 | 0.764 | 0.928 | 11th | 3th |
| 5 | *GCLM* | 0.823 | 0.700 | 0.946 | 23th | 2th |
| 6 | *IRS2* | 0.801 | 0.826 | 0.777 | 8th | 31th |
| 7 | *SESN2* | 0.801 | 1 | 0.602 | 1th | 80th |

## 4 Discusssion

This study demonstrated that the most powerful predictors of CR-relatedness across ageing-related genes rely on features heavily based on curated biological knowledge (from the literature), hereafter called ‘knowledge-based features’, such as GO terms and biological pathways. However, one caveat of these features is the difficulty to directly produce new findings, as they are based on existing knowledge. Nonetheless, the predictive power of these types of features enabled the extraction of additional biological insights from ageing-related genes strongly predicted to be CR-related but lacking current annotation of CR-relatedness.

Features not so heavily based on curated biological knowledge, particularly those based on gene expression and co-expression, were the least predictive, predicting almost randomly and implying that CR-relatedness is unlikely to be explained by gene expression analysis across tissues or by co-expression of ageing-related genes with other genes.

Upon merging the 9 datasets with specific feature types into two “whole” datasets (using two merging approaches), the predictive power was found to diminish compared to the strongest feature type-specific datasets. This occurred despite the use of a simple, univariate feature filtering algorithm. This suggests the importance of exploring the use of more sophisticated feature selection techniques, which is out of the scope of this work and left for future research.

The joint analysis of the top CR-candidate genes in {PathDIP, CAT} and {GO terms, BRF} models highlighted genes whose CR-relatedness can be explored from both GO terms and biological pathways perspectives. The strengths of both models were complementary, as {PathDIP, CAT} was more suited for classifying *Ageing*_*NotCR*_-related genes while {GO terms, BRF} performed better with *Ageing*_*CR*_-related genes. It is also noticeable that both models achieved similar predictive accuracies despite exhibiting only a moderate correlation of CR-related class predictions. In addition, top-ranked *Ageing*_*NotCR*_-related genes with high CR-probabilities were mostly surrounded by *Ageing*_*CR*_-related genes in the plane defined by the normalised CR-probabilities in {PathDIP, CAT} and {GO terms, BRF} models, suggesting biological process and pathway similarities between the top CR-related candidates and currently established CR-related genes.

### 4.1 Features importance

#### 4.1.1 GO term Features

The oxidation-reduction process was the most significant GO term for discriminating between the presence or absence of CR-relatedness across ageing-related genes. This term was also notable as it was associated with the greatest number of ageing-related genes across all top features in both PathDIP and GO term feature sets. Nonetheless, it did not demonstrate any significant preference for *Ageing*_*CR*_-nor AgeingNotCR-related genes, indicating that its effects on ageing could be mediated in both CR and NotCR-dependent ways. Evidence indicates that CR improves redox state (Kowaltowski, 2011), though this may not be the mechanism by which CR prolongs life (Lennicke and Cochemé, 2020). It has been observed, however, that low levels of ROS may actually be beneficial as mediators of redox signalling (Lennicke and Cochemé, 2020).

Notably, ageing-related genes associated with glucose and carbohydrate transport were almost exclusively *Ageing*_*CR*_-related, which at first sight may partially suggest a relationship between ageing, CR and glucose transport. Nevertheless, this relationship, if existing, is not straightforward as abnormal glucose metabolism is a common but not necessary feature of ageing (Kalyani and Egan, 2013). The most significant changes in glucose metabolism are due to ageing-related insulin dysfunction (Boemi et al., 2016). This phenomenon appears, however, to be a consequence rather than a cause of ageing, as the improvement in insulin sensitivity induced by CR was not required for the effects of CR on fitness and longevity (Dommerholt et al., 2018).

Ageing-related genes within the GPCR signalling pathway were also significantly related with CR. Some GPCRs have emerged as promising targets for reversing senescence and thus ageing (Santos-Otte et al., 2019). In this regard, one of the few studies discussing the relationship between a GPCR, namely *TGR5*, and CR (Wang et al., 2017) demonstrated that CR benefits on renal function ageing can be partially explained by up-regulation of *TGR5* ; as a result, the authors of that study proposed up-regulation of *TGR5* as a possible CR-mimetic candidate for renal function.

#### 4.1.2 PathDIP Features

The strongest predictive feature was autophagy, which is responsible for the disposal and recycling of metabolic macromolecules and damaged organelles (Chung and Chung, 2019). The second most significant feature, the longevity regulating pathway, is also associated with autophagy via a well-characterized signalling cascade (Chung and Chung, 2019). The fact that the *Ageing*_*CR*_-related genes in this study were significantly strongly associated with these autophagy-related pathways supports current hypotheses that some of the CR-related anti-aging effects are mediated by autophagy (Donati et al., 2008; Manco and Mingrone, 2005).

The BDNF pathway is presumably involved in brain ageing. Moreover, it is well established that CR enhances *BDNF* in a currently unknown manner (Manchishi et al., 2018; Budni et al., 2015; García-Prieto and Fernández-Alfonso, 2016), and that *BDNF* declines with age (Erickson et al., 2010). The possible relationship between this gene and CR could be explored through the well-characterized CR-related protein kinase B (Akt) pathway, as *BNDF* and *Akt* indirectly interact (Chen, 2019). However, in the context of our ML algorithms, it is possible that the BDNF pathway favoured NotCR-related predictions, as only one of its 11 ageing-related genes was predicted as CR-related.

*NRF2* is absent from GenDR. Nevertheless, our results suggest a strong association between the NRF2’s pathway and CR because, while the ratio of *Ageing*_*CR*_-related to *Ageing*_*NotCR*_-related genes is approximately 1/10 in the overall dataset, this pathway demonstrated the much greater 16/10 ratio; and this pathway has by far the greatest proportion of *Ageing*_*CR*_-related genes, compared to *Ageing*_*NotCR*_-related genes across all top PathDIP pathways. This finding could be supported by recent evidence linking *NRF2* and CR (Pomatto et al., 2020).

The “cellular responses to external stimuli” pathway is notable for the large number of ageing-related genes associated with it, only outnumbered, across all the most relevant features discussed in this work, by the “oxidation-reduction process” GO term. However, unlike the redox GO term, the relative distribution of ageing-related genes in this pathway did achieve statistical significance in direction of CR. External stimuli responses include responses to metal ions (Kultz, 2005), from where a metal ion theory of ageing was constructed. This theory has been poorly explored and opens opportunities for novel CR-research directions as it has been shown that CR decreases the level of certain metal ions in cells (Sharma et al., 2011).

### 4.2 CR-related candidate genes

*GOT1* and *GOT2* are genes whose products are involved in the amino acid metabolism that exist in cytoplasmic and mitochondrial forms, respectively (Stelzer et al., 2016). *GOT1* ‘s expression has been shown to change with age (Craig et al., 2015), but evidence linking it to CR is far scarcer. To our knowledge, only one study (Song et al., 2015) has demonstrated this relationship and proposed the role of *GOT1* as a significant metabolic feature associated with hepatic response to CR that is representative of differences in mediating amino acid influx into the gluconeogenic pathway. While there is lack of evidence linking *GOT2* with CR, one study (Miller et al., 2005) suggested that either *GOT1* or *GOT2* may impact H2S homeostasis, opening a window for further CR-related insights, as H2S signalling cascade has been observed to promote CR-like pro-longevity effects (Ng et al., 2018).

The TSC complex is a critical negative regulator of mTORC1 (Stelzer et al., 2016), the inhibition of which is associated with CR-like benefits (Madeo et al., 2019). In this regard, even if the TSC complex’s role in regulating mTORC1 *in vivo* remains under-explored, one study (Harputlugil et al., 2014) provides insights that indirectly link this gene with CR as it demonstrated that improved insulin sensitivity following short-term protein restriction (PR) required *TSC1* for facilitating increased pro-survival signalling after injury, and contributed to PR-mediated resistance to clinically significant hepatic ischemia reperfusion injury.

*CTH* is a gene that produces endogenous hydrogen sulphide (H2S) as a signalling molecule (Stelzer et al., 2016). Evidence linking *CTH* to CR is scarce. To our knowledge, only one recent study (Derous et al., 2017) reveals a positive correlation between *CTH* expression and CR application. This increased expression may have potential contributions to CR pro-longevity effects as inhibition of *CTH* is associated with about 15% lifespan reduction in worms. Moreover, *CTH* is a gene that promotes production of H2S, a potential CR-mimetic candidate, suggesting an approach for further studies linking *CTH* with CR.

The Glutamate-Cysteine Ligase Regulatory subunit (*GCLM*) is a gene that regulates the expression of antioxidant enzymes (Stelzer et al., 2016). Its role in CR is not explicitly stated in literature. However, a recent study (Lettieri-Barbato et al., 2020) showed its increased expression during fasting in *PASK* - deficient mice. Since fasting and intermittent fasting are associated to CR-like benefits, a link between *GCLM* and CR can be investigated from this perspective. If such a link does not exist, the outcome may remain informative by providing insights on differences between CR- and fasting-related beneficial signalling cascades.

IRS2 is a cytoplasmic signaling molecule that mediates the effects of insulin, insulin-like growth factor 1, and other cytokines. A homolog of this gene is present within GenDR (Gene Manipulations) (Wuttke et al., 2012), as “*chico*” in fruit fly, however, it was not detected as an ortholog by the OMA database (Train et al., 2017) and thus was not considered a CR-related gene. This gene has a further independent entry within GenDR (Gene Expression) where 174 mice genes that significantly change their expression during CR are reported (Wuttke et al., 2012; Plank et al., 2012). Out of our 7 top CR-candidate genes, *IRS2* was the only one overlapping these 174 genes as further explained in Supplementary materials (S.1.3). This suggests that *IRS2* does may not only be deferentially expressed during CR but also could have the potential to regulate CR-associated pro-longevity effects.

Sestrin 2 (SESN2) is an intracellular leucine sensor that negatively regulates the TORC1 signaling pathway. This gene was out of the scope of CR-relatedness until a recent work highlighted its role as a novel molecular link that mediates the effects of dietary amino acid restriction on TORC1 activity in stem cells of the fly gut, thereby maintaining gut health and ensuring longevity (Lu et al., 2021). Hence, although the CR-probability of this gene was relatively low in the {PathDIP, CAT} model, it was the highest in the {GO terms, BRF} model; and the averaged-based CR-related prediction of this gene is supported by recent evidence.

## 5 Conclusions

To our knowledge, this is one of the pioneering studies applying ML algorithms to CR research in the context of ageing. This work demonstrated the strong potential of ML-based techniques to identify CR-associated features as our findings are consistent with literature and recent discoveries. GO terms and PathDIP pathways were the most predictive types of features. Due to their curated knowledge-driven (literature-based) nature, the use of these feature types in the most predictive models has on one hand mostly corroborated existing knowledge (rather than directly generating new knowledge), but has on the other hand provided statistical support associating CR with the NRF2 pathway and GPCRs, which have been recently accumulating evidence towards CR in the literature, and so are worth further exploring.

Inference on novel CR-related features may be easier to accomplish from feature types not biased by curated knowledge. However, our work found an obstacle to this inference due to the low or even null predictive power of such feature types, implying that either 1) their features did not contain relevant information for predicting CR-relatedness; or 2) the used tree-based ensemble algorithms were not suitable for our classification problem with the unbiased feature types used in this work, especially expression and co-expression data; or 3) the number of currently known ageing-related genes was not large enough for our ML algorithms to find complex patterns in our unbiased data, leading to poor predictive accuracy in such datasets. In future work, the application of deep learning techniques could potentially increase the predictive power of unbiased feature types, which could provide novel insights on possible CR-related protein properties and interactions as well as CR-related gene expression and co-expression signatures.

Further insights were taken from genes annotated as *Ageing*_*NotCR*_-related genes in the dataset but strongly predicted as CR-related genes based on GO terms and PathDIP pathways. This analysis revealed a list of genes outside GenDR that are prone to be related with CR despite lacking such annotation. Most of these genes were consistent with some preliminary CR-related experiments, which makes them worth exploring for further wet-lab experiments to get a deeper understanding of their relationship with CR. Among these genes, *GOT2* was the only *Ageing*_*NotCR*_–related gene present within the top six stronger false positives in models learned with both PathDIP and GO term features. Other CR-related gene candidates strongly predicted by both most predictive ML models were *GOT1, TSC1, CTH, GCLM, IRS2* and *SENS2*, which, together with *GOT2*, remain to be validated in further lab-based experiments.

## Acknowledgments

This work was supported by a Biotechnology and Biological Sciences Research Council (BB/R014949/1) to J.P.M., a Wellcome Trust grant (208375/Z/17/Z) to J.P.M., a Leverhulme Trust research grant (RPG-2016-015) to J.P.M. and A.A.F., a CONACyT sponsorship (2019-000021-01EXTF-00468) to G.D.V.M, and an University of Guadalajara loan (V/2020/449) to G.D.V.M..

## References

Boemi, M., Furlan, G., Luconi, M.P., 2016. Molecular Basis of Nutrition and Aging: A Volume in the Molecular Nutrition Series. Academic Press, UOC Malattie Metaboliche e Diabetologia, INRCA-IRCCS, Ancona, Italy.

Budni, J., Bellettini-Santos, T., Mina, F., Garcez, M.L., Zugno, A.I., 2015. The involvement of bdnf, ngf and gdnf in aging and alzheimer’s disease. J Cell Mol Med 6, 331–341.

Carithers, L.J., Ardlie, K., Barcus, M., 2015. A novel approach to high-quality postmortem tissue procurement: The GTEx project. Biopreservation and Biobanking 13, 311–319. Analysis V8 database.

Chen, T., Guestrin, C., 2016. XGBoost: A scalable tree boosting system, in: Proceedings of the 22nd ACM SIGKDD International Conference on Knowledge Discovery and Data Mining, ACM, New York, NY, USA. pp. 785–794.

Chen, Y., 2019. Aging-induced akt activation involves in aging-related pathologies and aβ-induced toxicity. Aging Cell 18, e12989.

Chung, K.W., Chung, H.Y., 2019. The effects of calorie restriction on autophagy: Role on aging intervention. Ageing Research Reviews 11, 2923.

Craig, T., et al., 2015. The digital ageing atlas: integrating the diversity of agerelated changes into a unified resource. Nucleic Acids Res 43, D873–D878.

Csardi, G., Nepusz, T., 2006. The igraph software package for complex network research. InterJournal Complex Systems, 1695.

Dam, S.V., Craig, T., de Magalhaes, J.P., 2015. Genefriends: a human rnaseq-based gene and transcript co-expression database. Nucleic Acids Res 43, D1124–D1132.

Derous, D., et al., 2017. The effects of graded levels of calorie restriction: Xi. evaluation of the main hypotheses underpinning the life extension effects of cr using the hepatic transcriptome. Aging (Albany NY). 9, 1770–1804.

Dommerholt, M.B., Dionne, D.A., Hutchinson, D.F., Kruit, J.K., Johnson, J.D., 2018. Metabolic effects of short-term caloric restriction in mice with reduced insulin gene dosage. Redox Rep 237, 59–71.

Donati, A., Recchia, G., Cavallini, G., Bergamini, E., 2008. Relevance of autophagy induction by gastrointestinal hormones: Focus on the incretin-based drug target and glucagon. The Journals of Gerontology: Series A 63, 550–555.

Dorogush, A.V., Ershov, V., Yandex, A.G., 2018. Catboost: gradient boosting with categorical features support. 1706.09516 1.

Erickson, K.I., et al., 2010. Brain-derived neurotrophic factor is associated with age-related decline in hippocampal volume. J Cell Mol Med 30, 5368–5375.

Fabris, F., Freitas, A.A., 2016. New KEGG pathway-based interpretable features for classifying ageing-related mouse proteins. Bioinformatics 32, 2988–2995.

Fabris, F., de Magalhaes, J.P., Freitas, A.A., 2017. A review of supervised machine learning applied to ageing research. Biogerontology 18, 171–188.

Fabris, F., Palmer, D., Salama, K.M., de Magalhaes, J.P., Freitas, A.A., 2020. Using deep learning to associate human genes with age-related diseases. Bioinformatics 36, 2202–2208.

García-Prieto, C.F., Fernández-Alfonso, M.S., 2016. Caloric restriction as a strategy to improve vascular dysfunction in metabolic disorders. Circ Re 8, 370.

Gillespie, Z.E., Pickering, J., Eskiw, C.H., 2016. Better living through chemistry: Caloric restriction (CR) and CR mimetics alter genome function to promote increased health and lifespan. Frontiers in Genetics 7, 142.

Harputlugil, E., Hine, C., Vargas, D., Robertson, L., Manning, B.D., Mitchell, J.R., 2014. The tsc complex is required for the benefits of dietary protein restriction on stress resistance in vivo. Cell Reports 8, 1160–1170.

Huang, T., Zhang, J., Xu, Z.P., et al., 2012. Deciphering the effects of gene deletion on yeast longevity using network and machine learning approaches. Biochimie 94, 1017–1025.

Jalili, A., Salehzadeh-Yazdi, A., Asgari, Y., Arab, S.S., Yaghmaie, M., Ghavamzadeh, A., Alimoghaddam, K., 2015. Centiserver: A comprehensive resource, web-based application and R package for centrality analysis. PLoS ONE 10.

Jalili, M., Salehzadeh-Yazdi, A., Gupta, S., Wolkenhauer, O., Yaghmaie, M., Resendis-Antonio, O., Alimoghaddam, K., 2016. Evolution of centrality measurements for the detection of essential proteins in biological networks. Frontiers in Physiology 7, 375.

Kalyani, R.R., Egan, J.M., 2013. Diabetes and altered glucose metabolism with aging. Endocrinol Metab Clin North Am 42, 333–347.

Kowaltowski, A.J., 2011. Caloric restriction and redox state: Does this diet increase or decrease oxidant production? Redox Rep 16, 237–241.

Kultz, D., 2005. Molecular and evolutionary basis of the cellular stress response. Annu Rev Physiol 67, 225–57.

Lemaître, G., Nogueira, F., Aridas, C.K., 2017. Imbalanced-learn: A python toolbox to tackle the curse of imbalanced datasets in machine learning. Journal of Machine Learning Research 18, 1–5.

Lennicke, C., Cochemé, H.M., 2020. Redox signalling and ageing: insights from drosophila. Biochem Soc Trans 48, 367–377.

Lettieri-Barbato, D., Minopoli, G., Caggiano, R., Izzo, R., Santillo, M., Aquilano, K., Faraonio, R., 2020. Fasting drives nrf2-related antioxidant response in skeletal muscle. Int. J. Mol. Sci. 21, 7780.

López-Otín, C., Blasco, M.A., Partridge, L., Serrano, M., Kroemer, G., 2013. The hallmarks of aging. Cell 153, 1194–1217.

Lu, J., Temp, U., Müller-Hartmann, A., Esser, J., Grönke, S., Partridge, L., 2021. Sestrin is a key regulator of stem cell function and lifespan in response to dietary amino acids. Nature Aging 1, 60–72.

MacNee, W., 2016. Is chronic obstructive pulmonary disease an accelerated aging disease? Annals of the American Thoracic Society 13.

Madeo, F., Carmona-Gutierrez, D., Hofer, S.J., Kroemer, G., 2019. Caloric restriction mimetics against age-associated disease: Targets, mechanisms, and therapeutic potential. Cell Metabolism 29, 592–610.

de Magalhaes, J.P., Budovsky, A., Lehmann, G., Costa, J., Li, Y., Fraifeld, V., Church, G.M., 2009. The human ageing genomic resources: online databases and tools for biogerontologists. AgingCell 8, 65–72.

de Magalhães, J.P., Wuttke, D., Wood, S.H., Plank, M., Vora, C., 2012. Genome-environment interactions that modulate aging: powerful targets for drug discovery. Pharmacol Rev 64, 88–101.

Manchishi, S.M., Cui, R.J., Zou, X.H., Cheng, Z.Q., Li, B.j., 2018. Effect of caloric restriction on depression. J Cell Mol Med 22, 2528–2535.

Manco, M., Mingrone, G., 2005. Effects of weight loss and calorie restriction on carbohydrate metabolism. Curr Opin Clin Nutr Metab Care. 8, 431–9.

Menze, B.H., et al., 2009. A comparison of random forest and its gini importance with standard chemometric methods for the feature selection and classification of spectral data. BMC Bioinformatics 10.

Miller, R.A., Buehner, G., Chang, Y., Harper, J.M., Sigler, R., Smith-Wheelock, M., 2005. Methionine-deficient diet extends mouse lifespan, slows immune and lens aging, alters glucose, T4, IGF-I and insulin levels, and increases hepatocyte MIF levels and stress resistance. Aging Cell. 4, 119–25.

Ng, L.T., Gruber, J., Moore, P.K., 2018. Is there a role of H2S in mediating health span benefits of caloric restriction? Biochemical Pharmacology 149, 91–100.

Osorio, D., Rondon-Villarreal, P., Torres, R., 2015. Peptides: A package for data mining of antimicrobial peptides. The R Journal 7, 4–14.

Pedregosa, F., Varoquaux, G., Gramfort, A., Michel, V., Thirion, B., Grisel, O., Blondel, M., Prettenhofer, P., Weiss, R., Dubourg, V., Vanderplas, J., Passos, A., Cournapeau, D., Brucher, M., Perrot, M., Duchesnay, E., 2011. Scikitlearn: Machine learning in Python. Journal of Machine Learning Research 12, 2825–2830.

Plank, M., Wuttke, D., van Dam, S., Clarkeab, S.A., de Magalhaes, J.P., 2012. A meta-analysis of caloric restriction gene expression profiles to infer common signatures and regulatory mechanisms. Mil BioSyst 9, 1339–1349.

Pomatto, L.C.D., et al., 2020. Deletion of nrf2 shortens lifespan in c57bl6/j male mice but does not alter the health and survival benefits of caloric restriction. Free Radical Biology and Medicine 152, 650–658.

Rahmati, S., Abovsky, M., Pastrello, C., Jurisica, I., 2017. pathDIP: an annotated resource for known and predicted human gene-pathway associations and pathway enrichment analysis. Nucleic Acids Res 45, D419–D426.

Rainer, J., 2017. EnsDb.Hsapiens.v86: Ensembl based annotation package. R package version 2.99.0.

Rainer, J., Gatto, L., Weichenberger, C.X., 2019. Ensembldb: An R package to create and use ensembl-based annotation resources. Bioinformatics 35, 3151–3153.

Ri, J., Kim, H., 2020. G-mean based extreme learning machine for imbalance learning. Digital Signal Processing 98, 102637.

Ritchie, M.E., Phipson, B., Wu, D., Hu, Y., Law, C., Shi, W., Smyth, G., 2015. Limma powers differential expression analyses for RNA-sequencing and microarray studies. Nucleic Acids Res 43, e47.

Santos-Otte, P., et al., 2019. G protein-coupled receptor systems and their role in cellular senescence. Computational and Structural Biotechnology Journal 8, 1265–1277.

Sharma, P.K., Mittal, N., Deswal, S., Roy, N., 2011. Calorie restriction up-regulates iron and copper transport genes in saccharomyces cerevisiae. Mol Biosyst 7, 394–402.

Song, Y.M., et al., 2015. Metformin alleviates hepatosteatosis by restoring sirt1-mediated autophagy induction via an amp-activated protein kinase-independent pathway. Autophagy 11, 46–59.

Stark, C., et al., 2006. BioGRID: a general repository for interaction datasets. Nucleic acids research 34, D535–D539. Release 3.5.185): BIOGRID-MV-Physical-3.5.181.tab2.zip.

Stelzer, G., et al., 2016. The genecards suite: From gene data mining to disease genome sequence analysis. Current Protocols in Bioinformatics.

Tacutu, R., Thornton, D., Johnson, E., et al., 2018. Human ageing genomic resources: new and updated databases, build 20 (09/02/2020). Nucleic Acids Research 46, D1.

Train, C.M., Glover, N.M., Gonnet, G.H., Altenhoff, A.M., Dessimoz, C., 2017. Orthologous matrix (oma) algorithm 2.0: more robust to asymmetric evolutionary rates and more scalable hierarchical orthologous group inference. Bioinformatics 33, i75–i82.

Wang, X.X., et al., 2017. A dual agonist of farnesoid x receptor (fxr) and the g protein–coupled receptor tgr5, int-767, reverses age-related kidney disease in mice. Computational and Structural Biotechnology Journal 292, 12018– 12024.

Weidner, C.I., Lin, Q., Koch, C.M., 2014. Aging of blood can be tracked by DNA methylation changes at just three CpG sites. Genome Biology 15, R24.

Wieser, D., Papatheodorou, I., Ziehm, M., Thornton, J.M., 2011. Computational biology for aging. Philos Trans R Soc Lond B Biol Sci 366, 51–63.

Wuttke, D., Connor, R., Vora, R., et al., 2012. Dissecting the gene network of dietary restriction to identify evolutionarily conserved pathways and new functional genes, build 4 (24/06/2017). PLoS Genetics 8, e1002834.

Xiao, N., Cao, D.S., Zhu, M.F., Xu, Q.S., 2015. protr/protrweb: R package and web server for generating various numerical representation schemes of protein sequences. Bioinformatics 31, 1857–1859.

Zhavoronkov, A., Mamoshina, P., Vanhaelen, Q., Scheibye-Knudsen, M., Moskalev, A., Aliper, A., 2019. Artificial intelligence for aging and longevity research: Recent advances and perspectives. Ageing Research Reviews 49, 49–66.

