## supplementary material for "Machine-learning-based predictions of caloric restriction associations across ageing-related genes"

#### Supplementary texts

#### S.1 More on Datasets Constructions

##### S.1.1 KEGG-Pertinence Dataset construction

KEGG pathways can be regarded as directed graphs where each node $N$ is typically associated with either one gene product (*e.g.*, $N = \{G_1\}$) or more (*e.g.*, $N = \{G_1, G_2, G_3, ...G_n\}$), provided that the set of genes contained in a single node share similar functions and interactions. By contrast, edges represent how different nodes interact, mainly mediated by post-translational modifications and compounds.

The main difference between this perspective and the one that inspired it (Fabris and Freitas, 2016) is that here, the nodes in the pathways contain either one or multiple gene-products, as stated before; whereas, in (Fabris and Freitas, 2016), each node in the retrieved pathways was only associated with one single gene-product, which is why the authors used the term protein $p$, instead of node $N$, to refer to each one of the nodes in the pathways.

Since the creation of this dataset implied data extraction from each of the pathways' nodes and edges, the *KEGGlincs* package (White and Medvedovic, 2016a,b) was used to retrieve the KEGG Markup Language (KGML) representations of the 294 KEGG pathways obtained in the previously created *KEGG-Pertinence* dataset. These KGML representations consist of *igraph* objects (Csardi and Nepusz, 2006), where each *igraph* object contains the nodes and connectivity information of each of the pathways. Therefore, an instance or ageing-related reference node $N_{ref}$ in a KEGG pathway is a node that contains the ageing-related gene of interest, while the features for that instance are the remaining nodes in the pathway (which contains other gene-products that may be related to ageing, CR, or none of them) whose computation score is explained in the following paragraphs.

Feature values in this dataset quantify how each $N_{ref}$ influences all the other nodes in the pathway (*i.e.*, feature nodes) (Fabris and Freitas, 2016). The idea is that ageing-related nodes that have a common influence on a set of downstream nodes share a similar function. For each pathway, the influence score of $N_{ref}$ on a given node $N$ has the minimum value of 0 when the reference node $N_{ref}$ does not influence $N$ at all, because $N$ is not downstream of (*i.e.*, cannot be reached from) $N_{ref}$. On the other hand, the score for a given node $N$ that is downstream of $N_{ref}$ has the maximum value of 1 if, when $N_{ref}$ is removed from the pathway, the downstream node $N$ becomes unreachable from the nodes that are not downstream nodes of $N_{ref}$. In addition, if the score of a given node $N$ that is downstream of $N_{ref}$ has a value of 0.5, it means that $N_{ref}$ accounts for half of the influence that the downstream protein $N$ receives. A complete description of this calculation is provided in (Fabris and Freitas, 2016), where the concept of protein $p$ is used instead of node $N$.

The influence calculation is repeated for every $N_{ref}$ and $N$ nodes in every KEGG pathway retrieved for the KEGG-Pertinence dataset and the results were merged in such a way that, if a reference node $N_{ref}$ has different influence score values over $N$ for different pathways, the scores are summed, allowing for values greater than 1, which breaks the interpretation given in the last paragraph, where values lie within the interval [0,1]. In order to recover this interpretation, all the final scores are then divided by the total number of KEGG pathways used for the analysis (312). Thus, a final score for the pair $\{N_{ref}, N\}$ is the mean influence that $N_{ref}$ exerted on $N$ across all the pathways. The resulting dataset, which contained 1770 nodes as features and 799 ageing-related nodes as references, was called the KEGG-Influence dataset.

##### S.1.2 Protein descriptors

The features used for the protein descriptors dataset are as follows:

**Sequence Length and Molecular Weight** The sequence length is simply the count of amino acids of a protein. The molecular weight is the sum of the molecular weights of all amino acids in the protein.

**Amino Acid, Dipeptide, and Tripeptide Composition** It describes the fraction of each amino acid type as well as every possible dipeptide and tripeptide combinations within a protein sequence, giving rise to 20,400, and 8000 features, respectively.

**Z-values** The z-values, also known as Sandberg descriptors (Sandberg et al., 1998), are the principal components of 26 different physicochemical measured and calculated properties of amino acids, and essentially represent hydrophobicity/hydrophilicity ($z_1$), steric/ bulk properties and polarizability ($z_2$), polarity ($z_3$), and electronic effects ($z_4$ and $z_5$) of the amino acids.

**Local Descriptors** Local descriptors are computed based on the variation of functional groups defined by the occurrence of specific amino acids within the primary sequence of each protein (Silla and Freitas, 2011; Dubchak et al., 1995,

1999). These functional groups are defined based on the following physicochemical properties: Hidrophobicity, Normalised Van Der Waals volume, Polarity, Polarizability, Charge, Secondary structure and Solvent Accessibility. For each one of these properties, three groups are defined according to Table S1.

There are three types of local descriptors: Composition, Transition and Distribution:

- *Composition:* Accounts for the percentage composition (relative frequency) of a particular functional group within the amino acid sequence. Therefore, there are three composition features, one for each functional group of amino acids.

- *Transition:* Transition features represent the relative frequency in which an amino acid from a particular functional group is followed by an amino acid from another functional group. More precisely, the following transitions are considered: Group1 → Group2 or Group2 → Group1; Group1 → Group3 or Group3 → Group1; and Group2 → Group3 or Group3 → Group2.

- *Distribution:* Distribution features are computed based on the percentage of how many amino acids with a particular functional group are present on the first, 25%, 50%, 75% and 100% of the amino acid sequence.

In total, local descriptors in this dataset are composed of 21 features (3 compositions, 3 transitions, 15 distributions) for each of the seven properties, which was equivalent to 147 features. When adding the Z-values (5 features), amino acid composition (20 features), sequence length, and molecular weight (2); the final dataset contained 174 features.

#### S.2 More on Machine Learning Methods

##### S.2.1 Preprocessing

Two distinct preprocessing methods were applied, depending on the type of data to be computed. First, as explained in *S.2.1*, for binary features (*i.e.*, PathDIP, KEGG-Pertinence, PPI-Adjacency, GO terms) a minimum occurrence threshold was defined in order to filter features whose minority value of occurrence is equal or less than a grid of 3, 4 and 5 repetitions. On the other hand, continuous features (*i.e.*, KEGG-Influence, GTEx, and Proteins descriptors) were filtered based on a correlation criterion where one of each two features with correlation greater than 99% was eliminated.

The co-expression dataset was preprocessed differently from other continuous datasets due to its remarkably larger number of features that made correlation analysis highly demanding in terms of computational power and time. To address this, an F-statistic-based univariate feature filtering algorithm was applied using the *SelectKBest* command of the *sklearn.feature_selection* python's library to retrieve the 1000 genes that co-express the most with the CR-related target labels. Then, similarly to other continuous features, the 99% correlation-based filtering was applied to the remaining 1000 genes.

WholeDatasets' features were classified into two categories: binary, that underwent a 3-4-5 threshold preprocessing step; and continuous, to which the univariate filter was applied to reduce the number of continuous features to 1000, followed by the 99% correlation step. Some KEGG-Influence-based features had only two values and thus were preprocessed as binary within the WholeDatasets, even though this dataset was independently computed as continuous. This implies that some KEGG-Influence-based features within the WholeDatasets were preprocessed by the threshold criterion, while others by the univariate feature plus the correlation criterion.

##### S.2.2 Hyperparameters Grid

A brief description of the tuned hyperparameters of the classifiers is as follows:

- *Sampling Strategy:* It is the ratio of the number of instances associated with the minority class (*i.e.*, $Ageing_{CR}$-related genes) to the number of instances associated to the majority class (*i.e.*, $Ageing_{NotCR}$-related genes) after resampling. This is expressed as

$$\propto_{us}= \frac{N_m}{N_{rM}}, \tag{S.1}$$

  where $N_m$ is the number of instances in the minority class and $N_{rM}$ is the number of instances in the majority class after resampling.

- *Max. Features:* The number of randomly sampled features, from all $n$ features in the dataset, assigned as candidate features for selection in each node of a decision tree in the forest.

- *Num. Estimators:* The number of trees in the forest.

- *Class weights:* For each class label $y$ across the target labels, the *Balanced* mode computes a class weight $W_y$ that is proportionally inverse to the number of occurrences of $y$ within the training set as follows:

$$W_y = \frac{Num.\ Instances}{Num.\ Classes\ \times\ Occurrences\ of\ y}. \tag{S.2}$$

  The *Balanced_subsample* mode is the same as *Balanced* mode except that weights are computed based on the bootstrap sample for every tree grown. If this hyperparameter is set to *None*, all class labels are assigned the weight 1.

- *Binary Threshold:* It is the minimum number of occurrences that the minority value of a binary feature is required to have for that feature to be kept by a threshold-based filtering algorithm. This only applies to binary features.

Given the previous definitions, the following grid in Table S2 was established. It is to be noted that these parameters were not applicable or customisable to all classifiers. In case they do not, and for all the remaining customisable parameters provided by the ML libraries that do not appear in Table S2, default parameters were kept.

##### S.2.3 Analysis of GenDR (Gene expression) genes based on CR-association probability

We contrasted our estimated CR-association probabilities against pertinence to the GenDR (Gene Expression changes) database to see if our algorithm predicts ageing-related genes that undergo expression changes during CR. The following terminology is used:

- $CR_{GM}$-related genes: the 152 human orthologs genes retrieved from mice, fruit fly, nematodes and yeast in GenDR (Gene Manipulations). Manipulation of these genes is associated with modifications of CR-related pro-longevity effects (Wuttke et al., 2012). These genes are the "CR-related genes" reported in the main text.
- $CR_{GE}$-related genes: 143 human orthologs genes retrieved from the 174 mice genes with significant expression changes due to CR in GenDR (Gene Expression). Among the 143 human genes, 78 were up-regulated and 47 down-regulated. For mice, 101 were up-regulated and 73 down-regulated (Wuttke et al., 2012; Plank et al., 2012).
- $CR_{GM}$-probability: Probability of any given gene to be $CR_{GM}$-related according to the arithmetic mean of the of the computations given by {GO terms, BRF} and {PathDIP, CAT}. This is the same CR-probability reported in the main text.

We checked for the $CR_{GE}$-related genes that are also ageing-related (Figure S3) as their $CR_{GM}$-probability was already computed from the {GO terms, BRF} and {PathDIP, CAT} models. Only 18 out of 143 $CR_{GE}$-related genes overlapped with the ageing-related genes. The overlap between $CR_{GE}$-related with $CR_{GM}$-related genes is even smaller (9 genes). We then retained only 16 genes that had entries for both GO terms and PathDIP pathways.

Results (Table S3) indicate that only 7 ageing-related genes got a $CR_{GM}$-probability score equal or above 0.50 and thus were classified as $CR_{GM}$-related. IRS2 got the highest $CR_{CM}$-probability across the $CR_{GE}$-related genes. This gene is also one of the top $CR_{CM}$-related gene candidates reported in the main text and the only $CR_{GE}$-related gene that overlapped with the top $CR_{GM}$-candidates, suggesting that beyond being up-regulated during CR, manipulation of this gene may also be associated with changes on the ageing-related effects associated with CR. The following three top $CR_{GE}$-related genes classified as $CR_{GM}$-related (GCK, FMO3 and MAT1A) were actually $CR_{GM}$-related as they are already within the GenDR (Gene Manipulations) database. The remaining

3 of these 7 terms (GYS2, INMT, ARNLT) were just weakly $CR_{GM}$-predicted as they got a score equal or just above the 0.5 threshold.

### Supplementary Tables

Table S1: Classification of amino acids into three possible functional groups according to their corresponding physicochemical properties.

| Property number | Property name | Group 1 | Group 2 | Group 3 |
|---|---|---|---|---|
| Prop.1 | Hydrophobicity | Polar<br>R, K, E, D, Q, N | Neutral<br>G, A, S, T, P, H, Y | Hidrophobicity<br>C, L, V, I, M, F, W |
| Prop.2 | Normalised van der Waals Volume | 0-2.78<br>G, A, S, T, P, D, C | 2.95-4.0<br>N, V, E, Q, I, L | 4.03-8.08<br>M, H, K, F, R, Y, W |
| Prop.3 | Polarity | 4.9-6.2<br>L, I, F, W, C, M, V, Y | 8.0-9.2<br>P, A, T, G, S | 10.4-13.0<br>H, Q, R, K, N, E, D |
| Prop.4 | Polarizability | 0-1.08<br>G, A, S, D, T | 0.128-0.186<br>C, P, N, V, E, Q, I, L | 0.219-0.409<br>K, M, H, F, R, Y, W |
| Prop.5 | Charge | Positive<br>K, R | Neutral<br>A, N, C, Q, G, H, I, L, M, F, P, S, T, W, Y, V | Negative<br>D, E |
| Prop.6 | Secondary Structure | Helix<br>E, A, L, M, Q, K, R, H | Strand<br>V, I, Y, C, W, F, T | Coil<br>G, N, P, S, D |
| Prop.7 | Solvent Accessibility | Buried<br>A, L, F, C, G, I, V, W | Exposed<br>R, K, Q, E, N, D | Intermediate<br>M, S, P, T, H, Y |

Table S2: Ensemble algorithms' hyperparameters grid for sampling strategy, maximal number of features allowed in each base estimator, number of base estimators, class weight and binary threshold. The definition of $\alpha_{us}$ is as expressed in equation (S.1). The number $n$ in Max features is the same as the total number of features in the dataset of interest. The last column shows the algorithm(s) that use each of the hyperparameters mentioned in the first column.

| Hyperparameter | Values | Algorithm |
|---|---|---|
| Sampling strategy | $\alpha_{us} = 1$ | BRF, EEC |
| Max features | $\sqrt{n}, \quad log_2(n)$ | BRF, XGB |
| Num. estimators | 500 | BRF, EEC, CAT |
| Class weights | None, Balanced,<br>Balanced_subsample | BRF |
| Binary Threshold | $3, 4, 5$ | ALL |

Table S3: CR-probability analysis of the subset of ageing-related genes that are deferentially expressed during CR.

| Gene | GenAge | GenDR Genetic Manipulations | GenDR Expression changes | | Predicted $CR_{CM}$-probability |
|---|---|---|---|---|---|
| | | | Pertinence | Regulation | |
| IRS2 | ✓ | · | ✓ | Up | 0.80 |
| GCK | ✓ | ✓ | ✓ | Down | 0.75 |
| FMO3 | ✓ | ✓ | ✓ | Up | 0.64 |
| MAT1A | ✓ | ✓ | ✓ | Up | 0.57 |
| GYS2 | ✓ | · | ✓ | Up | 0.54 |
| INMT | ✓ | · | ✓ | Up | 0.50 |
| ARNTL | ✓ | · | ✓ | Down | 0.50 |
| GHR | ✓ | ✓ | ✓ | Down | 0.46 |
| PDIA3 | ✓ | · | ✓ | Down | 0.45 |
| PER2 | ✓ | · | ✓ | Up | 0.44 |
| CYP2F1 | ✓ | · | ✓ | Down | 0.37 |
| ACLY | ✓ | · | ✓ | Down | 0.36 |
| HSPA5 | ✓ | · | ✓ | Down | 0.31 |
| ALDH1A1 | ✓ | · | ✓ | Up | 0.31 |
| CYP2E1 | ✓ | · | ✓ | Up | 0.30 |
| DECR2 | ✓ | · | ✓ | Up | 0.27 |

### Supplementary Figures

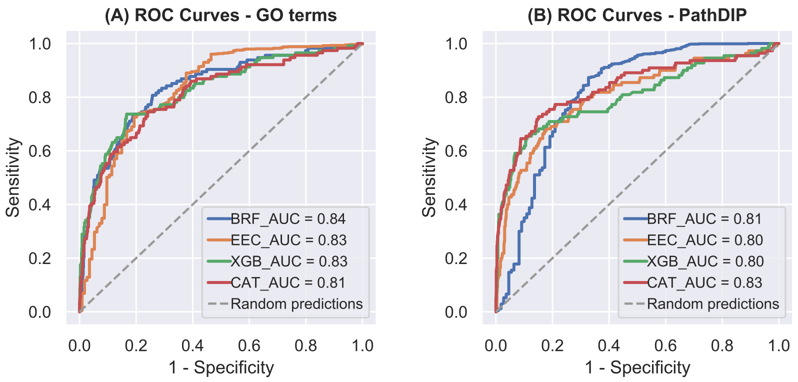


Figure S1: ROC curve and AUC of the ML algorithms on the two most predictive datasets.

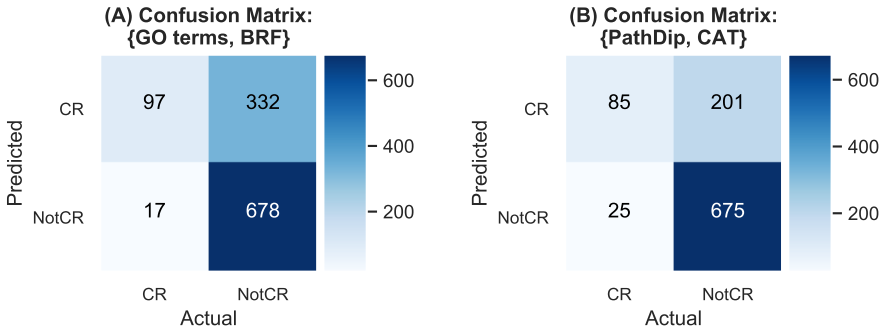


Figure S2: Comparison of the two most predictive models. **A** and **B**, confusion matrices of {GO terms, BRF} and {PathDIP, CAT}, respectively.

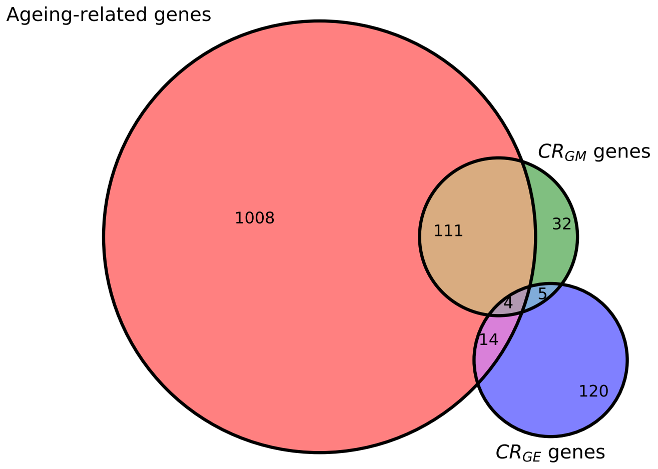


Figure S3: Overlapping between ageing-related, $CR_{GM}$-related and $CR_{GE}$-related genes.

#### References

Csardi, G., Nepusz, T., 2006. The igraph software package for complex network research. InterJournal Complex Systems, 1695.

Dubchak, I., Muchnik, I., Holbrook, S.R., Kim, S.H., 1995. Prediction of protein folding class using global description of amino acid sequence. Proceedings of the National Academy of Sciences 92, 8700–8704.

Dubchak, I., Muchnik, I., Mayor, C., Dralyuk, I., Kim, S., 1999. Recognition of a protein fold in the context of the scop classification. Proteins: Structure, Function and Genetics 35, 401–407.

Fabris, F., Freitas, A.A., 2016. New KEGG pathway-based interpretable features for classifying ageing-related mouse proteins. Bioinformatics 32, 2988–2995.

Plank, M., Wuttke, D., van Dam, S., Clarkeab, S.A., de Magalhaes, J.P., 2012. A meta-analysis of caloric restriction gene expression profiles to infer common signatures and regulatory mechanisms. Mil BioSyst 9, 1339–1349.

Sandberg, M., Eriksson, L., Jonsson, J., Sjostrom, M., Wold, S., 1998. New chemical descriptors relevant for the design of biologically active peptides. a multivariate characterization of 87 amino acids. J Med Chem 41, 2481–91.

Silla, C., Freitas, A.A., 2011. Selecting different protein representations and classification algorithms in hierarchical protein function prediction. Intelligent Data Analysis 15, 979–999.

White, S., Medvedovic, M., 2016a. KEGGlincs design and application: an R package for exploring relationships in biological pathways. URL: https://doi.org/10.7490/f1000research.1113436.1. version 1; not peer reviewed.

White, S., Medvedovic, M., 2016b. Visualize all edges within a KEGG pathway and overlay LINCS data. URL: http://www.bioconductor.org/packages/KEGGlincs. r package version 1.1.0.

Wuttke, D., Connor, R., Vora, R., et al., 2012. Dissecting the gene network of dietary restriction to identify evolutionarily conserved pathways and new functional genes, build 4 (24/06/2017). PLoS Genetics 8, e1002834.
